# Global molecular landscape of early MASLD progression in obesity

**DOI:** 10.1101/2025.01.13.632747

**Authors:** Qing Zhao, William De Nardo, Ruoyu Wang, Yi Zhong, Umur Keles, Gabriele Sakalauskaite, Li Na Zhao, Huiyi Tay, Sonia Youhanna, Mengchao Yan, Ye Xie, Youngrae Kim, Sungdong Lee, Rachel Liyu Lim, Guoshou Teo, Pradeep Narayanaswamy, Paul R. Burton, Volker M. Lauschke, Hyungwon Choi, Matthew J. Watt, Philipp Kaldis

## Abstract

Metabolic dysfunction-associated steatotic liver disease (MASLD) is often asymptomatic early on but can progress to irreversible conditions like cirrhosis. Due to limited access to human liver biopsies, systematic and integrative molecular resources remain scarce. In this study, we performed transcriptomic analyses on liver and metabolomic analyses on liver and plasma samples from morbidly obese individuals without liver pathology or at early-stage MASLD. While, the plasma metabolomic profile did not fully mirror liver histological features, dual-omics integration of liver samples revealed significantly remodeled lipid and amino acid metabolism pathways. Integrative network analysis uncoupled metabolic remodeling and gene expression as independent features of hepatic steatosis and fibrosis progression, respectively. Notably, GTPases and their regulators emerged as a novel class of genes linked to early liver fibrosis. This study offers a detailed molecular landscape of early MASLD in obesity and highlights potential targets of obesity-linked liver fibrosis.

**Graphical abstract:** 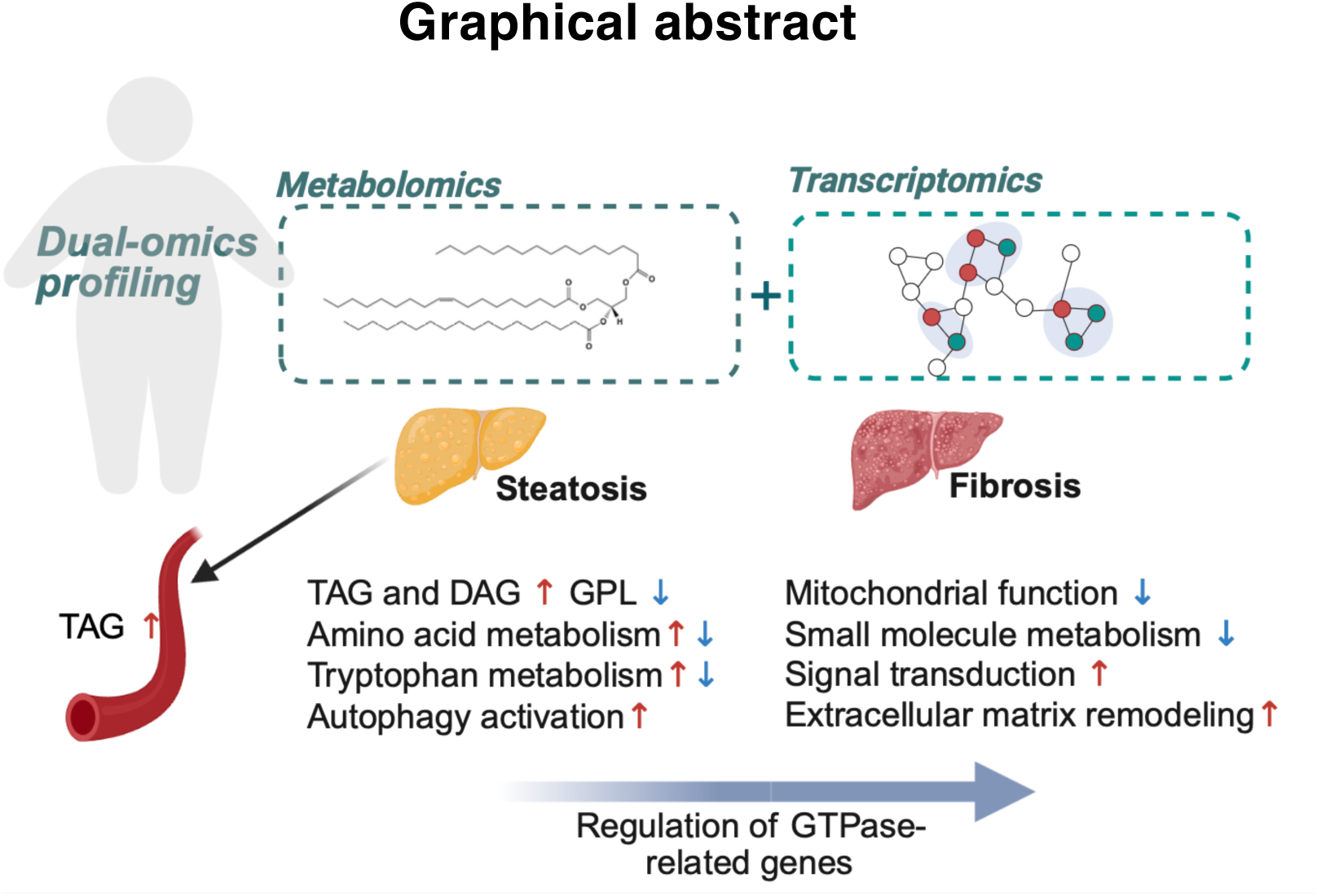

## Introduction

Metabolic dysfunction-associated steatotic liver disease (MASLD) is the most common chronic liver disease, affecting over 30% of adults worldwide (Miao et al., 2024). The prevalence of MASLD is closely associated with the epidemic of obesity, with an estimated 75% prevalence among individuals with obesity compared to 32% in the general population (Riazi et al., 2022). As liver manifestation of the metabolic syndrome, MASLD can cause, and is also caused by concurrent metabolic abnormalities commonly seen in obese individuals, such as insulin resistance, diabetes, dyslipidemia, and hypertension (Targher et al., 2024; Yki-Jarvinen, 2014).

Early stages of MASLD are characterized by hepatic steatosis, which is the excessive deposition of triacylglycerol (TAG) rich lipid droplets (LD) within the liver parenchyma and is usually reversible. However, in a subset of patients, the disease can develop into inflammatory and ballooning stages termed metabolic dysfunction-associated steatohepatitis (MASH), a more severe form of the disease with an increased incidence and severity of fibrosis. Fibrosis can be present at any stage of the MASLD spectrum of disease and at each stage is increasingly associated with treatment complications, liver-related and overall mortality (Angulo et al., 2015a; Ekstedt et al., 2015; Hagstrom et al., 2017). Unresolved fibrosis can progress to end-stage diseases including hepatic cirrhosis, and hepatocellular carcinoma, lethal malignancies with limited treatment options (Feng et al., 2024; Rodriguez et al., 2024).

Metabolic dysfunction is a primary feature and contributor of MASLD pathogenesis (Eslam et al., 2020). Hepatic steatosis is often accompanied by increased glucose production (Scoditti et al., 2024), elevated de novo lipogenesis (Lambert et al., 2014), and disrupted cholesterol homeostasis (Sakuma et al., 2025). Mitochondrial respiration is adaptively upregulated to prevent lipid accumulation in obesity (Koliaki et al., 2015), but with MASLD progression, mitochondria can become dysfunctional and exacerbate metabolic dysregulation (Fromenty & Roden, 2023). These interconnected metabolic phenotypes form the basis for the development of MASLD (Bril et al., 2017; Younossi et al., 2018). Recent advancements in metabolomics have greatly enhanced the understanding of the molecular systems biology of MASLD (McGlinchey et al., 2022; Vvedenskaya et al., 2021), however, few studies have combined such analysis with transcriptomics to provide integrative insights into disease pathogenesis in humans.

Fibrosis is the only histological feature of MASLD associated with liver-related mortality and morbidity (Angulo et al., 2015b). In the injured liver, damaged and apoptotic hepatocytes modulate the crosstalk between hepatocytes, liver macrophages (Kupffer cells), and hepatic stellate cells (HSCs) by releasing fibrogenic cytokines (e.g., TGF-β) and activating multiple signaling pathways (Bataller & Brenner, 2005; Subramanian et al., 2022). As a result of the multicellular response, HSCs transdifferentiate into active, collagen-producing myofibroblasts, driving fibrogenesis and the excessive deposition of extracellular matrix (ECM) (Subramanian et al., 2022). Since fibrosis is a key prognostic marker of MASLD progression, the shift from steatosis to fibrosis marks a critical point in the disease, offering a key opportunity for intervention to prevent further progression (Powell et al., 2021). Despite the availability of certain guidelines for clinical management of MASLD (European Association for the Study of the Liver. Electronic address et al., 2024; Rinella et al., 2023), the molecular and metabolic changes underpinning the key transitions driving disease progression in the context of obesity remain poorly understood. This poses a challenge to early disease management and prevention.

In this study, we generated a comprehensive map of gene expression and metabolomic profiles to delineate the molecular events associated with early-stage MASLD progression in obesity. Using this multi-omic resource, we focused our investigation on key molecular changes associated with (a) the transition from obesity with normal hepatic histology to MASLD and (b) the onset of liver fibrosis. Our data reveal distinct molecular signatures underlying steatosis and fibrosis progression, offering a detailed molecular portrait of the liver in early MASLD and highlighting potential therapeutic targets for reversing fibrosis at initial stages.

## Results

### Overview of the study

We analyzed samples from 109 obese individuals recruited before bariatric surgery at The Avenue, Cabrini, or Alfred Hospitals in Melbourne, Australia (**Table 1** and **Supplementary file 1**). Following the exclusion criteria described in Materials and Methods, 33 individuals lacked histological abnormalities (no MASLD) and 76 had MASLD (**Figure 1A**). Notably, 83 individuals (76%) were females in this cohort. Most individuals with obesity were in the early disease stages, with 74 individuals (68%) displaying grade 0-1 steatosis and 83 (76%) grade 0-1 fibrosis. Hepatic inflammation and ballooning were mild, with 9 cases exhibiting grade 2 or higher inflammation and only one case showing grade 2 ballooning (**Figure 1B** and **Table 1**). In clinical tests, MASLD patients displayed worse liver function (higher alanine transaminase (ALT)/aspartate aminotransferase (AST) ratio and gamma-glutamyl transpeptidase (GGT) levels), higher non-high-density (HDL) lipoprotein cholesterol and blood triglyceride levels, higher C-peptide, and insulin resistance compared to those with ‘No MASLD’ (**Table 1**, p < 0.05), confirming correct classification. Liver fibrosis was strongly correlated with insulin resistance, while steatosis grades were most correlated with the levels of plasma lipids (**Figure 1C**).

**Table 1.**
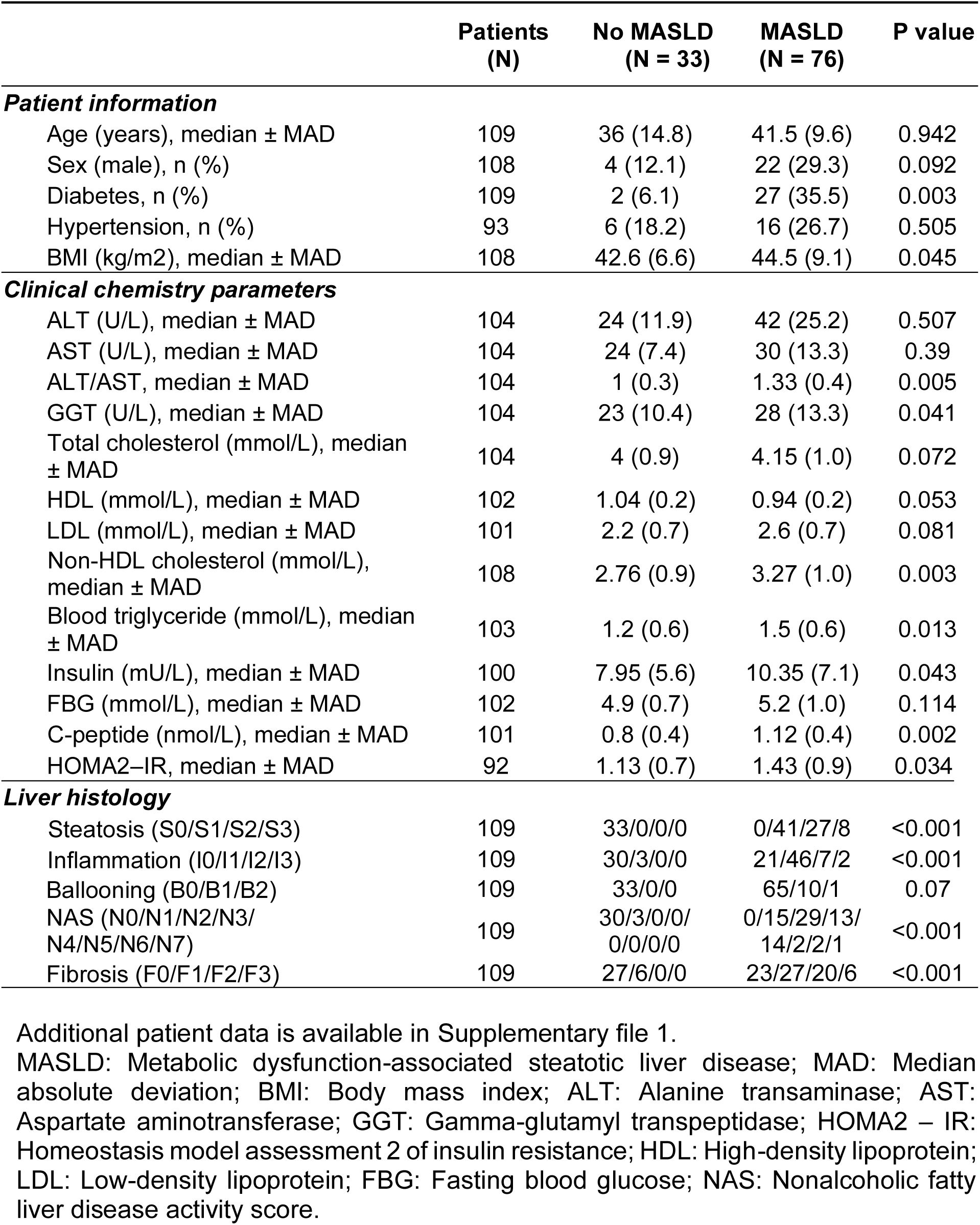
Patient characteristics.

|  | Patients<br>(N) | No MASLD<br>(N = 33) | MASLD<br>(N = 76) | P value |
| --- | --- | --- | --- | --- |
| **Patient information** |  |  |  |  |
| Age (years), median ± MAD | 109 | 36 (14.8) | 41.5 (9.6) | 0.942 |
| Sex (male), n (%) | 108 | 4 (12.1) | 22 (29.3) | 0.092 |
| Diabetes, n (%) | 109 | 2 (6.1) | 27 (35.5) | 0.003 |
| Hypertension, n (%) | 93 | 6 (18.2) | 16 (26.7) | 0.505 |
| BMI (kg/m<sup>2</sup>), median ± MAD | 108 | 42.6 (6.6) | 44.5 (9.1) | 0.045 |
| **Clinical chemistry parameters** |  |  |  |  |
| ALT (U/L), median ± MAD | 104 | 24 (11.9) | 42 (25.2) | 0.507 |
| AST (U/L), median ± MAD | 104 | 24 (7.4) | 30 (13.3) | 0.39 |
| ALT/AST, median ± MAD | 104 | 1 (0.3) | 1.33 (0.4) | 0.005 |
| GGT (U/L), median ± MAD | 104 | 23 (10.4) | 28 (13.3) | 0.041 |
| Total cholesterol (mmol/L), median ± MAD | 104 | 4 (0.9) | 4.15 (1.0) | 0.072 |
| HDL (mmol/L), median ± MAD | 102 | 1.04 (0.2) | 0.94 (0.2) | 0.053 |
| LDL (mmol/L), median ± MAD | 101 | 2.2 (0.7) | 2.6 (0.7) | 0.081 |
| Non-HDL cholesterol (mmol/L), median ± MAD | 108 | 2.76 (0.9) | 3.27 (1.0) | 0.003 |
| Blood triglyceride (mmol/L), median ± MAD | 103 | 1.2 (0.6) | 1.5 (0.6) | 0.013 |
| Insulin (mU/L), median ± MAD | 100 | 7.95 (5.6) | 10.35 (7.1) | 0.043 |
| FBG (mmol/L), median ± MAD | 102 | 4.9 (0.7) | 5.2 (1.0) | 0.114 |
| C-peptide (nmol/L), median ± MAD | 101 | 0.8 (0.4) | 1.12 (0.4) | 0.002 |
| HOMA2-IR, median ± MAD | 92 | 1.13 (0.7) | 1.43 (0.9) | 0.034 |
| **Liver histology** |  |  |  |  |
| Steatosis (S0/S1/S2/S3) | 109 | 33/0/0/0 | 0/41/27/8 | <0.001 |
| Inflammation (I0/I1/I2/I3) | 109 | 30/3/0/0 | 21/46/7/2 | <0.001 |
| Ballooning (B0/B1/B2) | 109 | 33/0/0 | 65/10/1 | 0.07 |
| NAS (N0/N1/N2/N3/N4/N5/N6/N7) | 109 | 30/3/0/0/0/0/0/0 | 0/15/29/13/14/2/2/1 | <0.001 |
| Fibrosis (F0/F1/F2/F3) | 109 | 27/6/0/0 | 23/27/20/6 | <0.001 |

Additional patient data is available in Supplementary file 1.
MASLD: Metabolic dysfunction-associated steatotic liver disease; MAD: Median absolute deviation; BMI: Body mass index; ALT: Alanine transaminase; AST: Aspartate aminotransferase; GGT: Gamma-glutamyl transpeptidase; HOMA2 – IR: Homeostasis model assessment 2 of insulin resistance; HDL: High-density lipoprotein; LDL: Low-density lipoprotein; FBG: Fasting blood glucose; NAS: Nonalcoholic fatty liver disease activity score.

**Figure 1.**
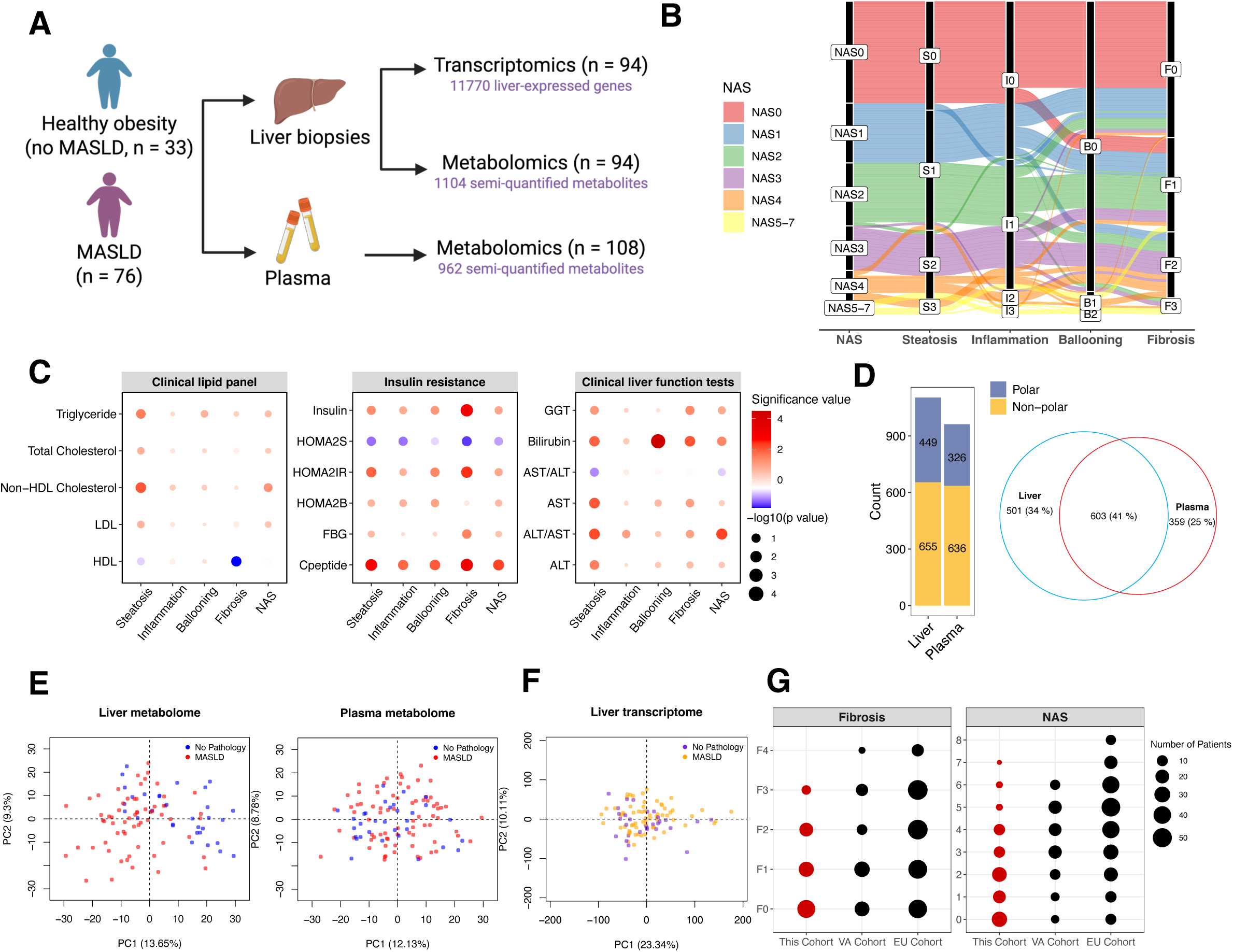
Study overview. (A) The overall study design. (B) Alluvial diagram of patient composition of groupings across liver histological features. (C) Relationship between clinical test results and stages of liver histological features. Node size was determined by –log10(p value) in ANOVA tests. Color indicates the significance degree [–log10(p value)] and the direction of change from early to late stages. (D) Number of metabolites analyzed in the liver and plasma. (E-F) Principal component analysis of liver metabolome, plasma metabolome, and liver transcriptome. (G) Disease spectra covered in this cohort and comparision to two published datasets.

We performed parallel transcriptomic profiling of liver slices and metabolomic analysis of liver and plasma samples (see Materials and Methods). Untargeted metabolomics (covering polar & non-polar metabolites including lipids) enabled semi-quantification of 1104 liver and 962 plasma metabolites, respectively (**Figure 1D**). Plasma urea and creatinine levels from untargeted metabolomics closely correlated with clinical assays (**Figure 2–figure supplement 1A**), supporting data reliability. Principal component analysis (PCA) revealed distinct clustering of the liver metabolome by MASLD status, but not in plasma (**Figure 1E**). The liver transcriptome was modestly separated between individuals with and without MASLD along the main principal components (**Figure 1F**), indicating that transcriptional programs may be shaped by both MASLD histological progression and confounding metabolic and biological processes in obese individuals (**Supplementary file 2**).

**Figure 2.**
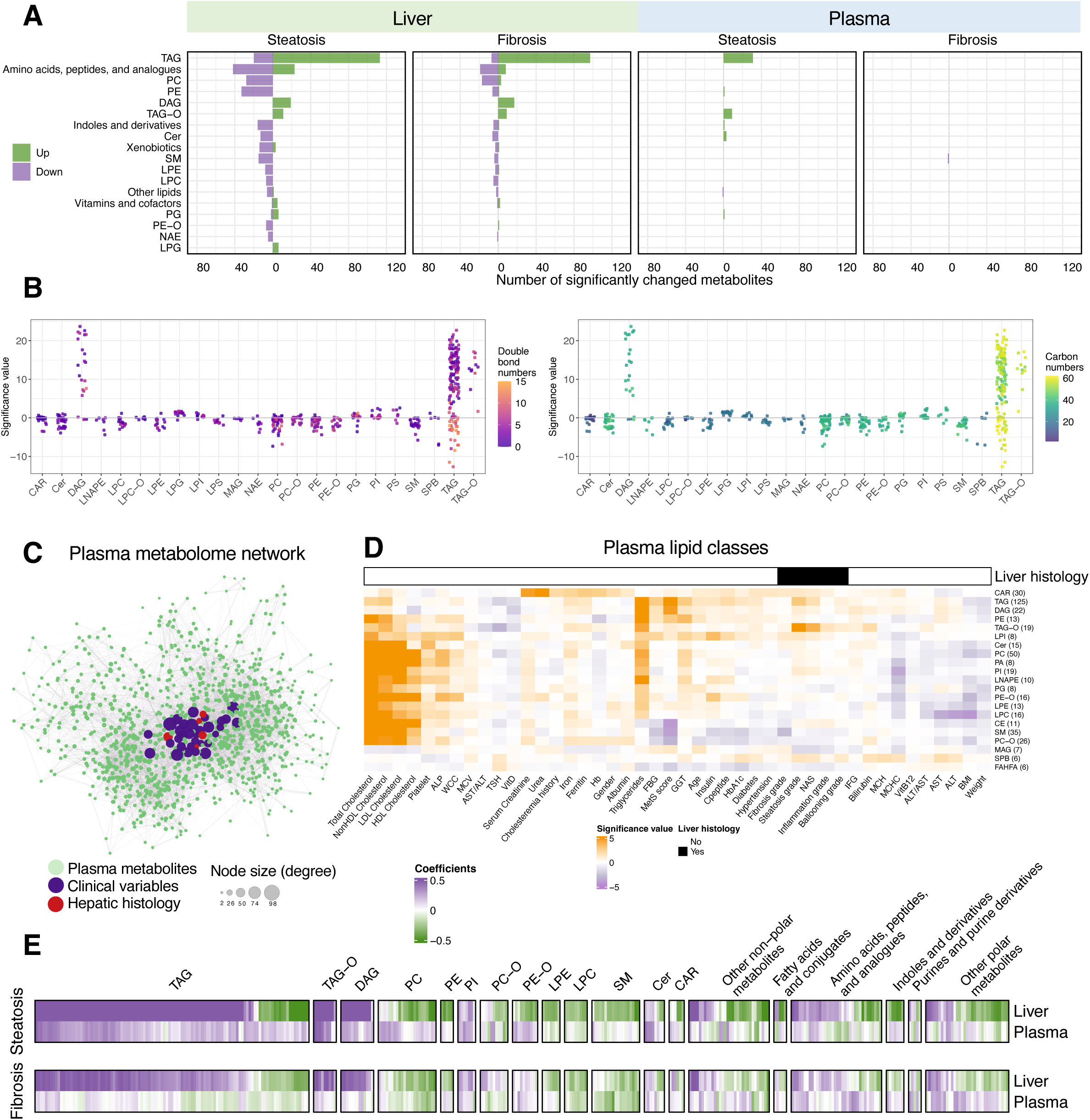
Hepatic and circulating metabolome in obese MASLD patients. (A) Number of metabolites in each class that were significantly associated with histological features in the liver (q value < 0.05). Metabolite classes with at least 5 metabolites with significance are shown in the plot. (B) Associations between steatosis grades and lipid species in each lipid class. Dots were colored by double bond numbers (left) and carbon numbers (right) of lipid species, respectively. (C) Partial correlation network of the plasma metabolome and clinical covariates. Node size reflects the degree of connectivity, with larger nodes indicating connections to a greater number of metabolite nodes. (D) Heatmaps for pairwise analysis between plasma lipid classes and clinical variables. Linear regression analysis was performed for numerical variables, while logistic regression was conducted for binary variables. Significance values refer to –log10(p value)*sign(coefficients) from regression models. (E) Comparisons of liver metabolome and plasma metabolome regarding their associations with liver steatosis and fibrosis.

As steatosis and fibrosis are the key histological features of MASLD, our statistical analyses focused on identifying molecular determinants linearly associated with their progression (**Supplementary file 3-5**). Models were adjusted for patient characteristics (age, sex, and BMI) and type 2 diabetes, the latter due to its differing prevalence across disease groups (**Table 1**). Moreover, we compared our transcriptomic data with those from two additional published cohorts: the Virginia cohort (VA cohort, GSE130970) (Hoang et al., 2019) with a disease spectrum comparable to that of our study, and the European cohort (EU cohort, GSE135251) (Govaere et al., 2020; Govaere et al., 2023) with a broader spectrum of the disease covering advanced MASLD pathologies (**Figure 1G**).

### Global investigation of the metabolome in obese MASLD patients

As expected, the liver metabolome was extensively remodeled in individuals with MASLD. Steatosis is the primary histological feature linked to liver metabolome, with 206 metabolites showing significant positive associations and 242 showing negative associations (q < 0.05, **Figure 2A** and **Figure 2–figure supplement 1B**). For example, we observed higher levels of glycerolipids (GLs, e.g. TAGs, diacylglycerols [DAGs]) and lower levels of membrane lipid classes, especially glycerophospholipids (GPLs) (**Figure 2A-B** and **Figure 2–figure supplement 1B**). Specifically, TAGs with 0 to 5 double bonds in the fatty acyl chains were positively associated with the histological outcomes, whereas TAGs containing at least one polyunsaturated fatty acid (PUFA) chain (e.g. number of double bonds > 5) were negatively associated with disease progression, particularly with steatosis (**Figure 2–figure supplement 1C**). Our findings are likely the results of increased de novo lipogenesis and elevated hepatotoxicity associated with saturated fatty acids (Roumans et al., 2020; Wang et al., 2006).

The plasma metabolome displayed limited associations with the histological features (**Figure 2A**), with statistically significant associations primarily with steatosis, such as TAGs (q value < 0.05, n = 22) and ether-linked TAG (TAG-O, q value < 0.05, n = 8) species. To better interpret the plasma metabolome data, we performed partial correlation network analysis to assess the associations among circulating metabolites, clinical variables, and hepatic histology in individuals with obesity (Lee et al., 2025). The network demonstrates the highest connectivity of plasma metabolites with clinical variables and fewer connections with hepatic histology features (**Figure 2C**). This implies that plasma metabolome largely reflects other metabolic conditions such as kidney function, dyslipidemia, and insulin resistance rather than hepatic features. Most plasma lipid classes were correlated with blood lipoprotein cholesterol levels (**Figure 2D**). However, in the context of obesity, blood lipoprotein levels did not show strong correlation with MASLD progression (**Figure 2–figure supplement 2**), indicating that their broad impact on plasma metabolome may mask the signals from hepatic abnormalities.

We next compared steatosis- and fibrosis-associated metabolite changes in the liver and plasma (**Figure 2E**). Plasma GLs exhibited similar but weaker associations with liver steatosis compared to hepatic GLs, whereas the depletion of PUFA-containing TAGs in the liver was not mirrored in circulation. Although plasma polar metabolites showed changes in the context of steatosis, potential circulating markers emerged for liver fibrosis with mild statistical significance. With fibrosis progression, there was a trend toward increased levels of tyrosine, quinolinic acid, lactic acid, and kynurenic acid in plasma (p < 0.05, q > 0.05) (**Supplementary file 3**), consistent with their reported roles as key hepatic metabolites implicated in MASLD progression and as proposed circulating biomarkers for MASLD severity or biopsy-proven MASH (Gou et al., 2025; Huang et al., 2025; Lahdou et al., 2013; Tan et al., 2025). Overall, despite elevated TAG levels and candidate fibrosis indicators, the circulating metabolome is less indicative of early-stage MASLD-related alterations in individuals with obesity and obesity-related co-morbidities.

### Integrative view of liver metabolism remodeling

To characterize metabolic remodeling in obese individuals during early disease development, we integrated the hepatic metabolome and transcriptome.

#### Lipid metabolism

In the liver, accumulation of GLs and mild reduction of GPLs were the main features of the metabolome changes (see **Figure 2**). Consistent with this observation, we identified genes implicated in the homeostasis of GLs and GPLs (**Figure 3A** and **Figure 3–figure supplement 1**). Genes such as *DGAT2*, *PNPLA3*, and *PLIN3* play a role in LD formation and TAG and DAG metabolism (**Figure 3–figure supplement 1B**) (Mashek, 2021). GPL metabolism was also markedly altered, including key genes such as *LPCAT1*, *PLD2*, *PCYT2*, *ETNK*2 and that are implicated in a compensatory response to the shift in phospholipid metabolism and the increased turnover of GPLs (Holdaway et al., 2025; van der Veen et al., 2012). In individuals with advanced hepatic fibrosis, expression levels of the genes involved in primary bile acid biosynthesis, such as *SLC27A5, AMACR*, *ACOX2*, *AKR1C4*, and *BAAT*, were lower. In addition, a group of cytochrome P450 (CYP) genes were downregulated during the progression of fibrosis, particularly those involved in CYP-dependent PUFA metabolism (i.e., linoleic acids and arachidonic acids into bioactive molecules) and steroid hormone biosynthesis (Dos Santos & Fleming, 2020; Hajeyah et al., 2020) (**Figure 3A**). Overall, our data highlights a number of genes encoding the enzyme subunits of anabolic and catabolic lipid metabolism in early MASLD, and some of these gene signals were consistently observed in both the EU and VA cohorts (Govaere et al., 2020; Hoang et al., 2019).

**Figure 3.**
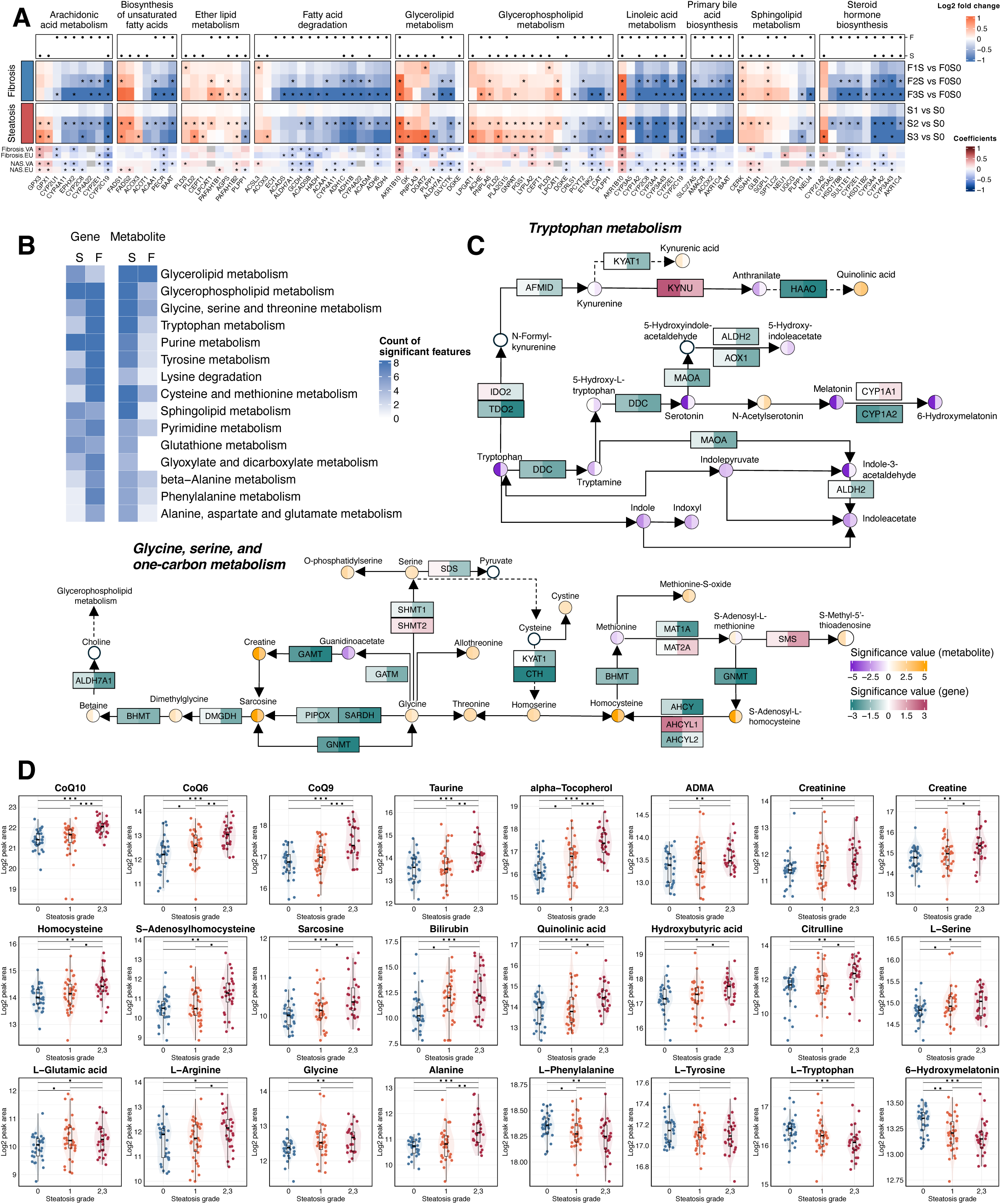
Integrative view of key metabolic pathways implicated in liver metabolism in obese individuals with MASLD. (A) Heatmap of log2 fold changes from pair-wise analysis of lipid metabolism-related genes. Association with steatosis (S) and fibrosis (F) is indicated by black dots (q value < 0.05, top). Results were cross-referenced with two published cohorts (VA cohort, GSE130970; EU cohort, GSE135251, bottom) Asterisks indicate genes with a q-value < 0.05 and a consistent change in direction within our cohort. (B) KEGG metabolic pathways with at least 4 genes or metabolites significantly associated with steatosis (S) or fibrosis (F). (C) Integrative map of gene and metabolite alterations associated with steatosis (left half of each box/circle) and fibrosis (right half of each box/circle). (D) Metabolite alterations corresponding to the advancement of steatosis grades (x-axis).

#### Dysregulated metabolic pathways during steatosis progression

To explore the variation in metabolic activities using dual-omic descriptors, we prioritized key pathways with both dysregulated metabolites and gene expressions (**Figure 3B**). In addition to significant GL and GPL remodeling, we also identified altered pathways of amino acid metabolism at both omics levels. Elevated levels of amino acids including glycine, glutamic acid, arginine, serine, and alanine in steatotic livers may reflect increased collagen synthesis and ECM remodeling during MASLD progression (Albaugh et al., 2017; de Paz-Lugo et al., 2018) (**Figure 3C-D**). Notably, aromatic amino acids including phenylalanine, tyrosine, and tryptophan were lower in subjects with advanced steatosis (**Figure 3D**). Dual-omics data revealed consistent downregulation of tryptophan metabolism, encompassing both the indole and melatonin pathways (**Figure 3C**). However, quinolinic acid, a key metabolite in the kynurenine pathway and a product of tryptophan catabolism, was significantly elevated in association with both hepatic steatosis and fibrosis. Collectively, early MASLD development is associated with reduced aromatic amino acid levels and the altered tryptophan catabolic flux in the liver, potentially reflecting the aberrant gut–liver axis communication in obese MASLD patients (Arto et al., 2024; Schnabl et al., 2025; Yanko et al., 2023).

Moreover, homocysteine and its upstream metabolite S-adenosylhomocysteine (SAH) levels were significantly higher in steatotic livers. The ratio of S-adenosylmethionine (SAM) to SAH and that of methionine to homocysteine indicated progressively decreasing trends with advancing steatosis (**Figure 3–figure supplement 2**), indicating impaired methylation potential under steatotic conditions (Walker, 2017). Furthermore, key regulators of SAM/SAH homeostasis, including *GNMT*, *MAT1A*, and *AHCY*, were downregulated in association with fibrosis (**Figure 3C**), which have been linked to liver pathogenesis in *in vivo* models (Alarcon-Vila et al., 2023; Matthews et al., 2009; Robinson et al., 2023; Varela-Rey et al., 2010). Collectively, these findings suggest that insufficient methylation modifications of DNAs, RNAs and proteins may be a mediator of MASLD pathogenesis, particularly in obesity.

In addition, we observed increasing levels of the antioxidants including coenzyme Q10 (CoQ10; ubiquinone), taurine, and alpha-tocopherol (vitamin E) with steatosis progression (**Figure 3D**). Yet another indicator of oxidative stress, hydroxybutyric acid, was increased in advanced steatosis and is also recognized as an early marker of insulin resistance (Sousa et al., 2021). Overall, the progression of liver steatosis is accompanied by extensive changes in amino acid metabolism and oxidative stress regulation.

#### Dysregulated genes involved in mitochondrial function and autophagy

In the liver metabolome, lower levels of hepatic long-chain acyl-carnitine (CAR) species were observed with advanced steatosis (CAR 18:2 and 20:4**, Supplementary file 4**), suggesting dysregulated fatty acid transmembrane transport and β-oxidation (Houten et al., 2016; Knottnerus et al., 2018). To systematically assess mitochondrial functions beyond β-oxidation, we mapped steatosis and fibrosis-associated genes to MitoCarta3.0 (Rath et al., 2021). Distinct mitochondrial dysfunction patterns emerged: steatosis involved 26 downregulated and 55 upregulated mitochondrial-function genes, including 16 related to mtDNA maintenance, while fibrosis was linked to 151 downregulated (dark green) and only 15 upregulated (orange) genes (**Figure 4A** and **Supplementary file 6**). These transcriptional shifts suggest that hepatic fibrosis involves broad mitochondrial impairment, contributing to oxidative stress and reduced ATP production, which could favor the exacerbation of steatohepatitis (Fromenty & Roden, 2023).

**Figure 4.**
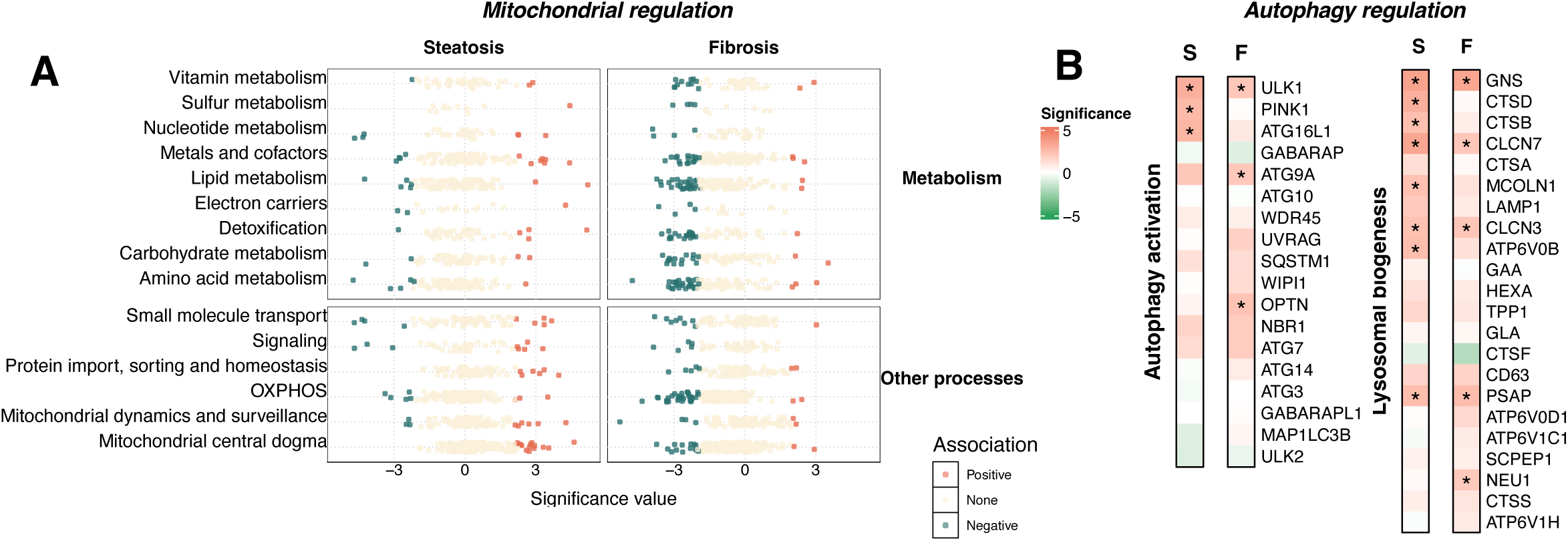
Dysregulated mitochondrial function and autophagy during the progression of steatosis and fibrosis. Gene expression patterns related to mitochondrial metabolism (A), mitochondria-related biological processes (A), autophagy activation (B), and lysosomal biogenesis (B) with statistical association to steatosis and fibrosis. S, steatosis. F, fibrosis.

Autophagy serves as a critical cellular response mechanism to the overload of intrahepatic TAG and cholesterol (Martinez-Lopez & Singh, 2015; Trivedi et al., 2021). Metabolomic analysis showed increased free cholesterol and reduced levels of CE 18:1 and 18:2, the predominant hepatic CE species (Xie et al., 2002) (**Figure 4–figure supplement 1A**), supporting prior findings that CE de-esterification contributes to elevated free cholesterol in MASLD (Min et al., 2012). Although autophagy has been linked to HSC activation (Kang & Chen, 2009; Tuohetahuntila et al., 2017), its involvement in MASLD progression remains unclear. Using a list of predictive genes for autophagy activation (Bordi et al., 2021), we observed upregulation of MTOR, a central member gene of mTORC1 complex that tightly regulates autophagy, with pro-autophagic markers such as *ULK1* and *PINK1* during steatosis progression (**Figure 4B**). Further investigation of over 600 autophagy genes (Bordi et al., 2021) (**Supplementary file 7** and **Figure 4–figure supplement 1B**) revealed active autophagy initiation in early MASLD, where 119 genes across pathways including mTORC and upstream effectors, lysosome, and autophagy core assembly genes displayed altered expression levels. This suggests that autophagy serves as an adaptive response to hepatic lipotoxicity during steatosis, but its activation may decline as the disease progresses, as evidenced by the downregulation of key regulators such as *NR1H4*, *SIRT5*, *FOS*, and *EGR1* (**Figure 4–figure supplement 1B**), thereby impairing liver metabolism (Singh et al., 2009).

### Molecular signatures of hepatic steatosis and fibrosis are mutually independent

To gain insights into the dual-omic data in the liver further, we performed a partial-correlation network analysis to integrate liver transcriptomic, metabolomic, and clinical data (Lee et al., 2025). The resulting subnetwork highlighted distinct molecular signatures correlated with steatosis and fibrosis (**Figure 5A**). Steatosis was primarily associated with hepatic neutral lipid accumulation and related metabolomic alterations, whereas fibrosis was predominantly linked to transcriptional changes, including the upregulation of genes involved in ECM remodeling (e.g., *TGFB3*, *FSTL1*), signal transduction (e.g., *RHOU*, *DOK3*), and metabolism (e.g., *CYP2C19*, *PNPLA4*, *SGPL1*). These findings underscore the presence of distinct molecular pathways driving the progression of steatosis and fibrosis.

**Figure 5.**
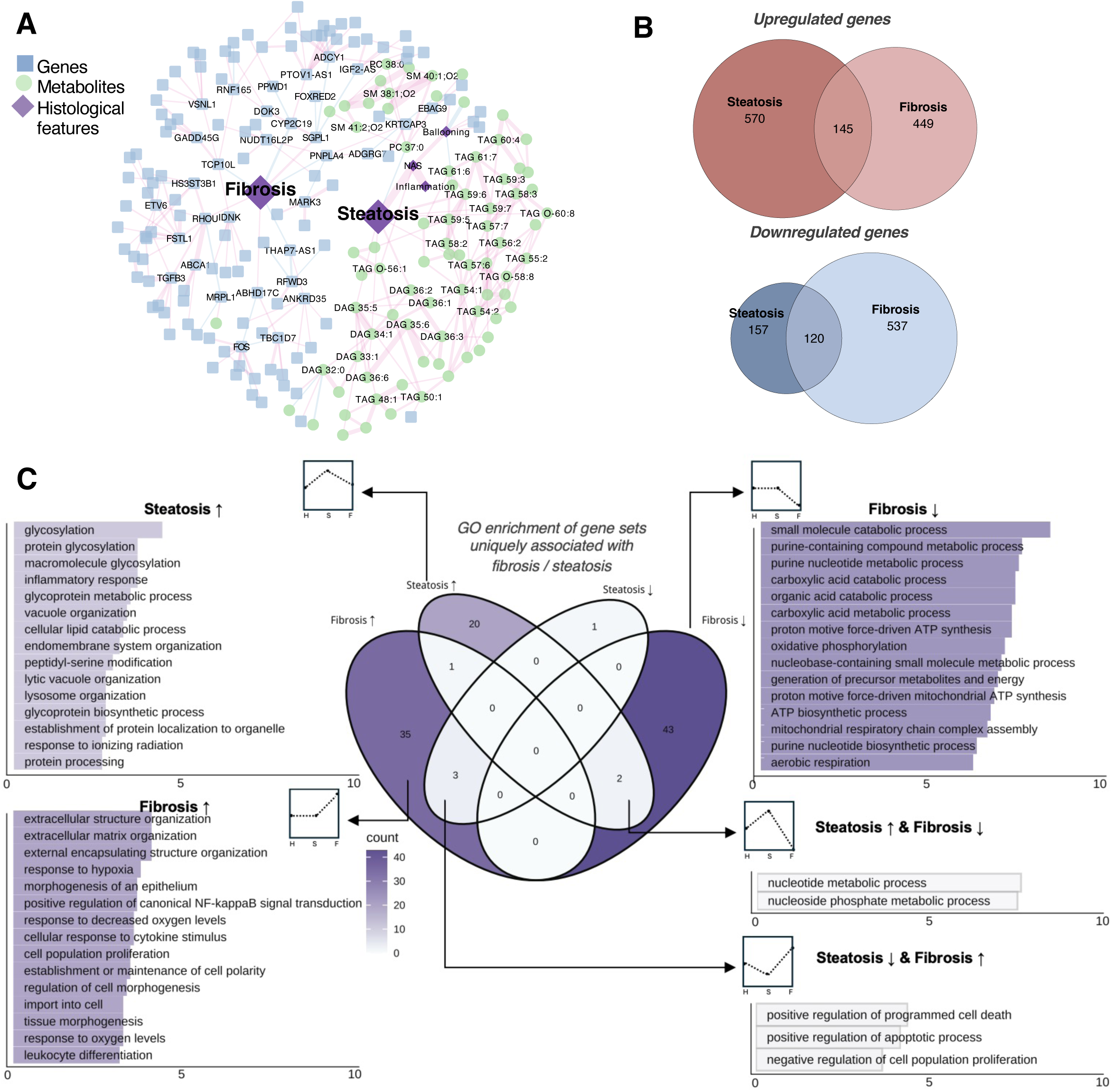
Steatosis and fibrosis as independent processes. (A) Network of genes, metabolites, and histological features (fibrosis and steatosis) using partial correlation analysis as described in the methods. (B) Venn diagrams depicting the number of genes significantly associated with steatosis and fibrosis (q value < 0.05). (C) Gene ontology (GO) enrichment of gene sets specific to steatosis or fibrosis. H, healthy obese controls. S, steatosis. F, fibrosis.

In our transcriptomics analysis, 992 and 1,251 gene markers were identified as significantly associated with the histological features of steatosis and fibrosis, respectively, with limited overlap between the two sets (**Figure 5B** and **Supplementary file 5**, q value < 0.05), suggesting unique transcriptomic regulations underlie each feature. Functional enrichment analysis further supported this distinction, revealing largely non-overlapping biological processes associated with each gene set (**Figure 5–figure supplement 1** and **Supplementary file 8**).

To distinguish gene signatures uniquely linked to steatosis or fibrosis, we included the other histological feature as a covariate in the regression model. This allowed us to define steatosis-specific genes (independent of fibrosis) and fibrosis-specific genes (independent of steatosis) (**Supplementary file 9**). Since steatosis precedes fibrosis in the pathophysiology of MASLD, enrichment analysis of steatosis-and fibrosis-specific genes revealed a ‘pseudo-temporal’ progression of biological processes (**Figure 5C**). More pathways were found to be associated with fibrosis than with steatosis. Steatosis-specific upregulated genes were enriched in protein glycosylation, inflammatory response, lipid catabolism, and lysosome organization. In contrast, fibrosis-specific genes showed downregulation of various metabolic pathways and upregulation of processes related to ECM remodeling, hypoxia, signaling, and cell morphogenesis. Additionally, apoptosis-related pathways were suppressed in steatosis but activated in fibrosis, while nucleotide metabolism showed the opposite trend, suggesting dynamic regulation of cell death and proliferation during MASLD development. These findings implicate distinct and lesion-specific gene regulatory programs in early MASLD progression.

Moreover, consistent with previous observations showing that hepatic fibrosis correlates with insulin resistance parameters in clinical assays (**Figure 1C**), we found that individuals with diabetic MASLD exhibited a greater number of downregulated genes as fibrosis progressed than non-diabetic MASLD individuals (**Supplementary file 10 and Figure 5–figure supplement 2**). Since most of the suppressed genes in the diabetic subgroup are involved in metabolism (e.g., *BAAT*, *G6PC1*, *SULT2A1*, *MAT1A*), we hypothesize that diabetes may exacerbate the metabolic dysfunction associated with hepatic fibrosis progression.

### Gene signatures of fibrosis initiation

To further explore representative gene signatures in the development of fibrosis, we identified 213 genes as progressive markers by comparisons of gene expression levels against two different baselines (no MASLD and steatosis; details provided in the methods; **Supplementary file 11**). Among them, 75 of these markers overlapped with fibrosis-associated genes in the VA cohort and 35 overlapped with the EU cohort (**Figure 6A**) (Hoang et al., 2019), resulting in 130 novel fibrosis markers. Pathway analysis of these fibrosis markers highlighted prominent roles for signal transduction and ECM organization/ disassembly, as expected (**Figure 6–figure supplement 1A**). To infer the cell type-of-origin for signals from bulk sequencing, we mapped progressive fibrosis markers onto a human liver single-cell atlas (MacParland et al., 2018). Upregulated genes appeared to reflect changes from different cell types, whereas downregulated markers were predominantly hepatocyte-specific (**Figure 6A**, left panel). This suggests that in the early stages of liver fibrosis development, the transcriptional programs are activated in different cell types, while hepatocyte functions potentially become muted during the process. Meanwhile, we observed fibrosis-associated upregulation of genes involved in TGF-β and SMAD signaling cascades (*TGFB1*, *TGFB3*, and *INHBA*), ECM activators (*ITGAV*, *LOXL2*, *THBS1*, *MMP9*, and *MMP14*) (Li et al., 2022; Munger et al., 1999; Wen et al., 2018), regulators (*CDK8*, *CTDSP2*, *HDAC1*, and *HDAC7*), and downstream targets (*TIMP1*, *MMP9*, and *COL5A2*) (**Figure 6–figure supplement 1B**). This aligns well with previous reports that TGF-β signaling and HSC activation drive fibrosis during MASLD progression (Kisseleva & Brenner, 2021; Massague & Sheppard, 2023).

**Figure 6.**
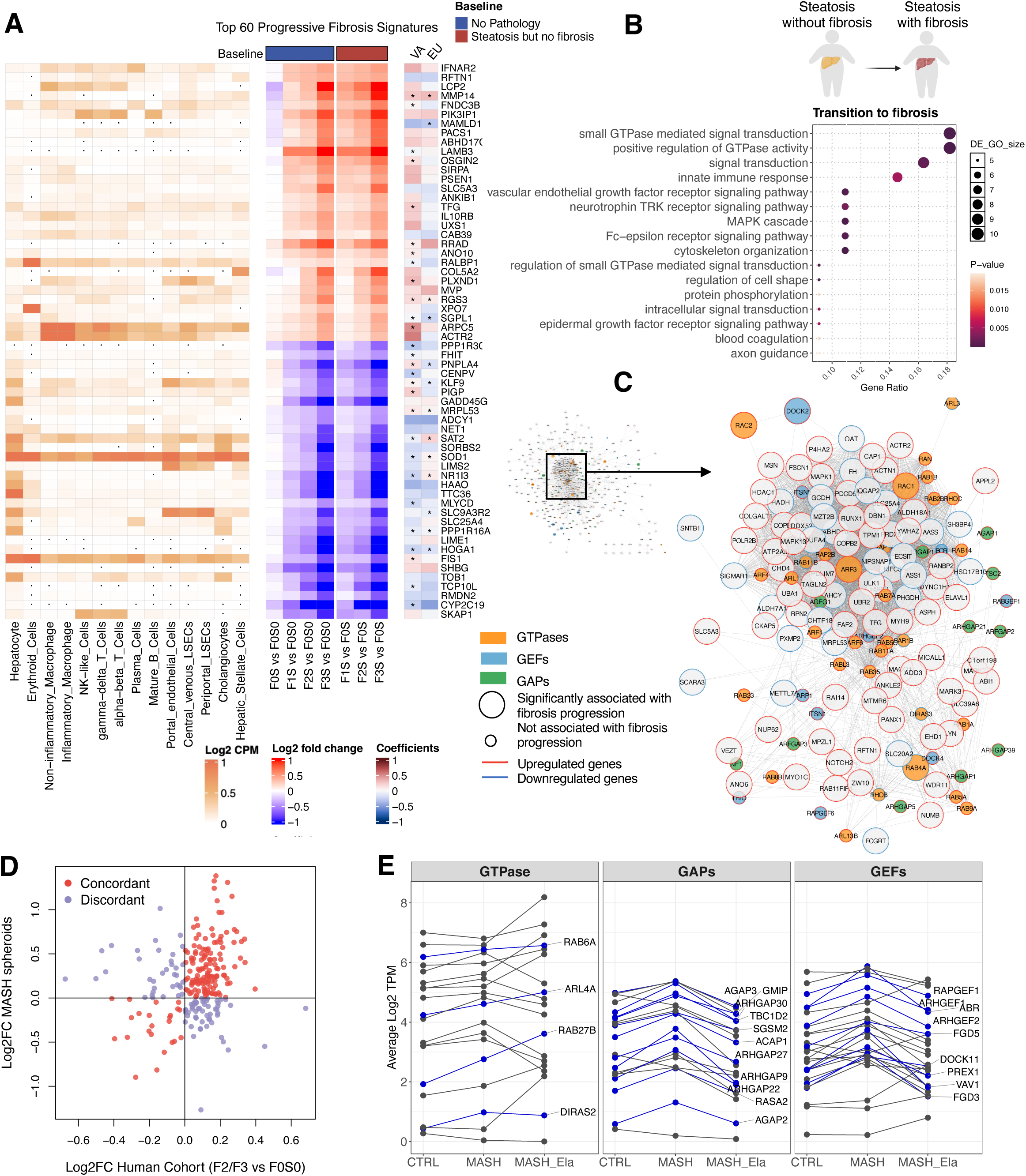
Liver fibrosis signatures and a potential therapeutic target of fibrosis initiation. (A) Top 60 fibrosis markers in the liver. A total of 213 genes were identified, from which the top 30 upregulated and top 30 downregulated genes are visualized. The left side of the plot (blue bar on top) shows the comparison of individuals with fibrosis and individuals without pathology (baseline - “no MASLD”). For the right side (red bar), individuals with fibrosis were compared to those with steatosis but no fibrosis (baseline – “steatosis but no fibrosis”). Results were cross-referenced with two published cohorts (VA cohort, GSE130970; EU cohort, GSE135251, right). Asterisks indicate genes with a q-value < 0.05 and a consistent change in direction within our cohort. The average gene expressions of liver cell types were obtained from GSE115469 at the log2CPM level. Dots in the single cell map indicate zero expression in the corresponding cell types. (B) Enrichment of pathways in the transition from simple steatosis to the onset of fibrosis. (C) Protein interaction network of GTPases and their regulators. Nodes are colored based on gene type, with borders indicating the direction of gene regulation. Node size corresponds to the significance of the genes in relation to fibrosis grades within this cohort. GEFs: Guanine nucleotide exchange factors. GAPs: GTPase-activating proteins. (D) Comparison of expression level changes in GTPase-related genes between this human cohort and an independent 3D spheroid MASH system. Log2 fold change for the human cohort was calculated by comparing patients with grade 2 or 3 fibrosis to those without fibrosis or steatosis. Genes with the same direction of change are colored in red, while others are colored in purple. (E) Expression of GTPase-related genes in patient-derived 3D liver spheroids (controls; MASH; MASH treated with Elafibranor; n = 4). Sixty-eight genes with a p-value < 0.05 from the ANOVA test were plotted in the diagram, with blue lines highlighting 24 genes that exhibited increased expression in the MASH group compared to the control group.

Given that the transition from simple steatosis to the onset of fibrosis marks a critical window for disease management, we compared gene expression profiles between individuals with fibrosis accompanied by steatosis and those with steatosis but without fibrosis (**Supplementary file 12**). Notably, the top-enriched pathways in this comparison emerged as GTPase signaling and its regulation, signal transduction, and innate immune response (**Figure 6B**) among other pathways. Moreover, we identified 37 GTPase-related genes displaying significant associations with fibrosis progression (**Figure 6–figure supplement 2A** and **Supplementary file 13**). To explore the potential role of GTPase signaling in fibrosis, we examined an inventory of 251 genes encoding GTPases, GTPase-activating proteins (GAPs), and guanine nucleotide exchange factors (GEFs), analyzing their interactome using the Human Cell Map (Go et al., 2021) and a protein-protein interaction network (Huttlin et al., 2017; Razick et al., 2008).

Among the 6,971 proteins that have been tested for physical interaction or subcellular proximity with GTPase-related genes, 508 of these associated significantly with fibrosis progression in our cohort (**Figure 6C**). The network indicates that GTPases and their regulators interact with a wide range of fibrosis genes, with *ARF3* and *RAC1* being central hubs of the network. Functional enrichment of the core nodes revealed that the GTPase network regulates intracellular transport, actin cytoskeleton organization, exocytosis, and other biological processes during fibrosis progression (**Figure 6–figure supplement 2B**). Therefore, GTPases and their regulators are co-regulated with fibrosis-related genes encoding their protein interaction partners, supporting the likelihood of a functional link between GTPases and hepatic fibrosis. This is supported by TGF-β genes positively correlated with co-expressed GTPase-related genes (**Figure 6–figure supplement 2C**), suggesting a potential link between TGF-β signaling and GTPase pathways.

To experimentally validate our findings, the altered expression of GTPase-related genes was explored in an independent system using an established model of 3D liver spheroids in which hepatic cells remain viable and functional for multiple weeks (Bell et al., 2016; Kemas et al., 2021; Vorrink et al., 2017). Specifically, we co-cultured primary fully-differentiated human hepatocytes with Kupffer and stellate cells isolated from adult patients with histologically confirmed MASH (Youhanna et al., 2025) and examined the overall expression profile of GTPase-regulated genes between our human cohort and the ex vivo system. The expression of GTPases were largely concordant when assessing human patients and the 3D spheroid system (**Figure 6D**). In comparison to the control group, spheroids from MASH patients demonstrated an upregulation of 24 genes encoding GTPases and their regulators (**Figure 6E**). Remarkably, the treatment of liver spheroids with elafibranor, a dual PPARα/δ agonist (Ratziu et al., 2016), restored the expression levels of 17 (70%) GTPase-related genes back to the baseline (**Figure 6E**). This independent experimental system further suggests that GTPases may play a role in MASLD pathogenesis and the fibrotic response.

Since HSCs are the main liver cell type responsible for activating fibrosis, we next assessed whether inhibition of GTPase activity in an immortalized HSC cell line, the LX-2 cells, could attenuate markers of fibrogenesis (**Figure 6–figure supplement 3A**). Selective GTPase inhibitors targeting Rac1 (NSC23766) and Cdc42 (ML141) reduced the mRNA expression of *COL1A1* and *COL1A2*, as well as pro-collagen secretion (**Figure 6–figure supplement 3**) under basal conditions. TGF-β1 mediated activation induced the expected increase in collagen secretion and gene expression which was attenuated by Rac1 inhibitor (NSC23766) and to some extent by ML141. In addition, examination of previously published transcriptomic data of HSCs isolated from CCL_4_-mediated liver fibrosis in mice (De Smet et al., 2021), revealed the upregulation of GTPases comparable to the steatosis-to-fibrosis transition in our human cohort, with a temporal pattern aligning with hepatic collagen deposition (**Figure 6–figure supplement 4A**). Further, in a human liver organoid model, TGF-β induced increased expression of GTPase-related genes in hepatocytes and HSCs, but not in fibroblasts (Hess et al., 2023) (**Figure 6–figure supplement 4B**), suggesting a potential feedforward loop involving the TGF-β/GTPase axis between hepatocytes and HSCs. To investigate intercellular crosstalk in GTPase regulation, we examined key GTPase-related genes in LX-2 cells, hepatocyte monocultures, and spheroid co-cultures (including hepatocytes, HSCs, Kupffer cells). As shown in **Figure 6–figure supplement 5**, TGF-β1 induced a potential increase in *VAV1* and *DOCK2* consistent in co-cultures, hepatocytes, and LX-2 cells, while *RAC1*, *RAB32*, and *RHOU* showed cell type-specific responses. These findings indicate that multiple hepatic cell types mediate GTPase regulation, underscoring intercellular crosstalk, which requires further detailed investigation.

Overall, these additional experiments and analyses of datasets identified upregulated GTPase-related genes during fibrosis initiation (**Figure 6–figure supplement 4C**). However, further in-depth mechanistic studies are needed to validate this association to determine how TGF-β is regulating GTPases and how GTPase control the secretion of collagen, leading to fibrosis.

## Discussion

In this study, we performed a comprehensive omics-based analysis of a cohort of 109 obese individuals with early MASLD. Through integrative dual-omics approaches, we mapped the liver and plasma metabolomes and identified distinct hepatic molecular features associated with steatosis and fibrosis progression. Notably, fibrosis was closely connected to global reprogramming of hepatic gene expression, with GTPase-related genes emerging as possible mediators of fibrosis initiation.

The strength of our study is the characterization of hepatic pathophysiology in relation to steatosis and fibrosis via transcriptome and metabolome-wide variations. Consistent with general expectation, our work highlighted altered hepatic lipid metabolism in human liver tissues of obese patients at both metabolite and gene levels (Liu et al., 2023; Radosavljevic et al., 2024; Saliba-Gustafsson et al., 2024). Consistent with the variations in GLs, higher expression levels of *DGAT2*, *PNPLA3*, and *PLIN3* were associated with steatosis progression. The higher expression of *DGAT2*, which encodes a key enzyme catalyzing the last step of de novo TAG synthesis, implies enhanced integration of TAGs into LDs in the endoplasmic reticulum, which presumably alleviates lipid-induced ER stress and evades accumulation of lipotoxic lipids (Scorletti & Carr, 2022). Upregulation of *PLIN3* was correlated with higher grades of both fibrosis and steatosis. The protein encoded by *PLIN3*, along with other PLIN proteins (Van Woerkom et al., 2024), plays an important role in LD stabilization and the prevention of TAG hydrolysis, thus potentially contributing to hepatic steatosis (Carr et al., 2012). In addition, although previous animal studies have reported increased levels of sphingolipids in MASLD models (Babiy et al., 2023; McGlinchey et al., 2022), we did not observe this in patients with obesity at early stages of the disease. Sphingolipid alterations varied across cohorts with differing patient compositions (McGlinchey et al., 2022; Ooi et al., 2021; Vvedenskaya et al., 2021), suggesting that sphingolipid metabolism may be dependent on disease stage, obesity status, and exhibit discrepancies between humans and mice.

Beyond lipid metabolism, other metabolic pathways and processes also exhibited dysregulation at the dual-omics level. Specifically, with excessive lipid deposition, we observed higher levels of taurine and vitamin E, both of which have antioxidant effects and likely reflect the hepatic response to oxidative stress (Al-Baiaty et al., 2021; Arroyave-Ospina et al., 2021; Wei et al., 2024). Importantly, amino acid metabolism was markedly dysregulated as steatosis progressed, characterized by elevated hepatic levels of serine, arginine, glutamic acid, glycine, and alanine, along with decreased levels of the aromatic amino acids phenylalanine, tryptophan, and tyrosine. A previous *in vitro* study also demonstrated that palmitic acid supplementation disrupted the metabolism of the aromatic amino acids in PH5CH8 and HepG2 cells (Aggarwal et al., 2023). As microbiota-derived metabolites, dysregulation in these amino acid may reflect disrupted gut function and may affect the de novo synthesis of nicotinamide adenine dinucleotide in MASLD patients (Xue et al., 2023). Moreover, we also observed evidence of autophagy activation concurrent with intrahepatic lipid and cholesterol accumulation, wherein lysosome-associated genes were uniquely associated with steatosis, as shown in **Figure 4B** and highlighted in **Figure 5C** by the enrichment of lysosome organization among steatosis-specific processes. This autophagic response most likely represents a protective mechanism to ameliorate steatosis (Gual et al., 2017); however, the downregulation of autophagy regulators during fibrosis development may exacerbate liver metabolic dysfunction.

As liver fibrosis progressed, the expression levels of genes involved in primary bile acid biosynthesis, PUFA metabolism, and steroid hormone biosynthesis were lower, especially *ACADS*, *ACADSB*, and *ACADM* which encode key enzymes in the initial stage of catalyzing fatty acid β-oxidation, and *HADH* and *ACAA1*, which encode enzymes that act in the late stage of β-oxidation (Adeva-Andany et al., 2019). Moreover, mitochondrial-related genes were downregulated with fibrosis progression, particularly those related to the electron transport chain of the mitochondrial inner membrane. This may indicate that pathological oxidative stress and impaired ATP production caused by early mitochondrial dysfunction may aggravate fatty liver inflammation (Koliaki et al., 2015), and the augmentation of the inflammatory response may become an important cause of liver fibrosis.

Through univariate hypothesis testing and partial correlation analysis, we revealed distinct molecular signatures correlated with fibrosis and steatosis. Liver fibrosis was associated with gene expression alteration, from which we identified over 200 progressive markers tracking its gradual advancement. Furthermore, GTPase-related genes were dysregulated at the mRNA level with the emergence and progression of fibrosis. Increased expression of GTPase regulators was independently confirmed in 3D primary human liver spheroids, with elafibranor restoring their expression to baseline levels (**Figure 6E**). The downregulated expression of GTPase regulators following elafibranor treatment suggests a potential anti-fibrotic mechanism involving GTPase signaling.

GTPase proteins are categorized into several subfamilies such as Rho-, Ras-, and Arf-GTPase based on their sequence and structure (Gray et al., 2020). GAPs and GEFs are the key regulators of GTPase activity that modulate intrinsic GTPase functions and the GTP-bound state (Bos et al., 2007). Only a few studies have elucidated the role of GTPases in liver pathology (Agarwal et al., 2023; Huang et al., 2018; Peng et al., 2021; Schwerbel et al., 2020). For instance, *Rap1a* was identified as a signaling molecule that suppresses both gluconeogenesis and hepatic steatosis (Agarwal et al., 2023). The Rab23-specific GAP, *GP73*, has been implicated in triggering non-obese MASLD (Peng et al., 2021). Additionally, Rho-GTPase signaling through the ROCK1/AMPK axis regulates de novo lipogenesis during overnutrition (Huang et al., 2018), while an immune-related GTPase triggers lipophagy and prevents hepatic lipid storage (Schwerbel et al., 2020). However, evidence regarding the role of GTPase regulation in human liver is limited, particularly in the context of liver fibrosis initiation and progression, and a thorough mechanistic investigation is warranted to establish causality.

In recent years, dual- and multi-omics strategies have gradually emerged in MASLD research. A proteo-transcriptomic map constructed from the EU cohort’s liver RNA sequencing data and circulating proteome identified four promising protein biomarkers (Govaere et al., 2023). Among these, *AKR1B10* was also identified in our study as a key gene related to lipid metabolism, showing strong associations with both steatosis and fibrosis (**Figure 3A**), which highlights its potential as a biomarker even in the early stages of MASLD. Additionally, we observed mitochondrial dysregulation linked to the progression of liver fibrosis (**Figure 4A**). Consistent with our findings, a multi-omics study in obese individuals with MASLD also revealed mitochondrial dysfunction at the hepatic protein level. The study further emphasized similar mitochondrial disruptions in adipose tissue, suggesting the liver-adipose tissue interaction in obesity-related MASLD (Castane et al., 2025). Beyond inter-tissue interactions, Raverdy *et al*. reported the heterogeneity of MASLD molecular signatures in relation to varying degrees of cardiovascular risk and type 2 diabetes (Raverdy et al., 2024). This further highlights the impact of systemic metabolic conditions on MASLD progression. In our analysis of diabetic individuals, we identified more downregulated metabolic genes associated with fibrosis, supporting the notion that insulin resistance may exacerbate metabolic dysregulation and interferes with the disease trajectory.

Overall, our integrative investigation offers one of the most detailed molecular landscapes of early stage MASLD in individuals with obesity, with comprehensive descriptors of the early remodeling of hepatic metabolism associated with initiation of fibrosis in the steatotic liver.

### Limitations of the study

This study focused on 109 patients with early-stage MASLD, potentially overlooking molecular changes associated with later disease stages. To address this, we cross-referenced our findings with two external cohorts (EU and VA). Moreover, as the results were based predominantly on participants without diabetes, their validity in diabetic populations warrants additional evaluation since a potential association between liver fibrosis and insulin resistance was observed. However, because of the limited number of diabetic cases and the imbalanced distribution of fibrosis grades between diabetic and non-diabetic groups, the interaction between fibrosis and diabetes could not be evaluated in this study.

We identified GTPase-related genes as potential future targets for reversing liver fibrosis. While preliminary validation has been conducted in different in vitro systems, further investigation is required to confirm the causal role of specific GTPases in fibrosis initiation especially in HSCs and to elucidate how GTPase signaling contributes to collagen production throughout the progression of MASLD.

We used patient derived liver spheroids as an independent experimental system to validate some of our data (see Figure 6D-E). This system has been extensively validated (Bell et al., 2018; Bell et al., 2016; Bell et al., 2017; Messner et al., 2018; Vorrink et al., 2017) and consists of adult patient-derived hepatocytes, hepatic stellate cells, and Kupffer cells that are fully differentiated. These spheroids are stable for several weeks in culture but as any in vitro culture system also has disadvantages. Given that the spheroids consists of 3 cell types, we cannot determine which cells are mainly driving the expression of the genes we are measuring. This is demonstrated in **Figure 6–figure supplement 5,** where some genes display higher expression in the Co-culture, in hepatocytes (Mono-culture), or LX-2 cells, respectively. To understand better how these cell types signal to each other, additional experiments will need to be performed.

## Materials and Methods

### Ethics statement

Participants provided written and verbal informed consent. The study protocol conforms to the ethical guidelines of the 1975 Declaration of Helsinki and was approved by the University of Melbourne Human Ethics Committee (ethics ID 1851533), The Avenue Hospital Human Research Ethics Committee (Ramsay Health; ethics ID WD00006, HREC reference number 249), the Alfred Hospital Human Research Ethics Committee (ethics ID GO00005), and Cabrini Hospital Human Research Ethics Committees (ethics ID 09-31-08-15). All work on the data from human samples in Sweden was approved by Etikprövningsmyndigheten (Dnr 2024-07917-01).

### Cohort recruitment

A detailed medical history was taken, and metabolic comorbidities were noted including the presence of previously diagnosed hypertension and diabetes assessed by oral glucose tolerance testing. Exclusion criteria included: age <18 years, previous gender reassignment, other causes of chronic liver disease and/or hepatic steatosis including Wilson’s disease, α-1-antitrypsin deficiency, viral hepatitis, human immunodeficiency virus, primary biliary cholangitis, autoimmune hepatitis, genetic iron overload, hypo- or hyperthyroidism, coeliac disease, as well as recent (within three months of screening visit) or concomitant use of agents known to cause hepatic steatosis including corticosteroids, amiodarone, methotrexate, tamoxifen, valproic acid and/or high dose oestrogens (Chalasani et al., 2017). Further exclusion criteria included potential for alcohol-induced liver disease, which was assessed through a modified version of the alcohol use disorders identification test (AUDIT) (Chalasani et al., 2017; Saunders et al., 1993). Following histological assessment, patients with liver slice weighing less than 2mg were excluded from the omics analysis.

Eligible patients with obesity scheduled for primary or secondary sleeve gastrectomy, gastric bypass, or the insertion of a laparoscopic-adjustable band were prospectively enrolled. All patients were fasted for 8-12 h overnight and venous blood was taken before the induction of anaesthesia. Blood was transferred to 2 x K2E Ethylenediaminetetraacetic acid (EDTA), 2 x SST™ II Advance and 1 x FX 5 mg bio-containers for subsequent storage or clinical/biochemical assessments. All blood samples were sent to Melbourne PathologyTM (Victoria, Australia) for standardised measurement of biochemical and metabolic variables, except for one bio-container of EDTA. Standard blood analyses were performed for electrolytes, full blood examination, glucose, glycosylated haemoglobin, insulin, C-peptide, cholesterol, triacylglycerols, and liver function assessed by ALT, AST, GGT, and alkaline phosphatase (ALP), and screening blood tests for liver disease. The remaining blood within the EDTA tube was spun at 8000 x g for 10 min and the plasma was collected and stored at −80°C for mass spectrometry analyses.

### Liver biopsy collection and histological feature assessment

An ∼ 1 cm^3^ wedge liver biopsy was collected from the left lobe of the liver during surgery. All liver samples were collected between 8am and 1pm. The liver was cut into two portions. One portion was placed in formalin and transported to TissuPath^TM^ (Mount Waverley, Victoria), paraffin embedded and processed for histological analysis. Samples were graded according to the Clinical Research Network NAFLD activity score (NAS) (Brunt et al., 2011) and Kleiner classification of liver fibrosis (Kleiner et al., 2005) by a liver pathologist at TissuPath. Patients with no MASLD were defined by steatosis score of 0, regardless of inflammation or fibrosis grade 1. MASL patients were defined by a steatosis score ≥1 with or without lobular inflammation. The remaining portion of liver slices was used for bulk RNA sequencing and untargeted metabolomics analysis.

### Sample preparation for untargeted metabolomics

For liver tissue biopsies, each sample containing 50 ± 5 mg of tissue was extracted in 0.8 mL 50:50 (v/v) methanol:chloroform (Wu et al., 2019; Xu et al., 2016). Samples were homogenized for 10 min at 25 Hz with a single 3-mm tungsten carbide bead per tube. Separation of phases was achieved by the addition of 0.4 mL of water followed by vortex-mixing and centrifugation (2400 g, 15 min, 4 °C). After separation, the upper phase (the metabolite-containing fraction) and the lower phase (the lipid-containing fraction) were transferred into separate tubes and dried using SpeedVac.

For plasma samples, 225 µL of methanol was added to 50 µl plasma, and the mixture was vortexed for 10 sec (Lee et al., 2014). Subsequently, 450 µL chloroform was added and the mixture was incubated for 1h in a shaker. To induce phase separation, 187.5 µL water was added and the mixture was centrifuged at 12,000 rcf for 15 min at 4 °C. The upper phase (the metabolite-containing fraction) phase and lower (the lipid-containing fraction) were collected into separate fresh tubes and dried using SpeedVac.

Dry extracts from the organic fraction were resuspended in the mixed solvent of 90:10 isopropanol: acetonitrile. For the aqueous fraction, 90% acetonitrile was used to reconstitute dry extracts. Plasma samples were reconstituted in 1:4 dilution (plasma-to-solvent: 50 μL/200 μL), and the reconstitution volumes for liver biopsy samples were corrected by the weights of liver slices (tissue-to-solvent: 50 mg/1000 μL). After reconstitution, samples were sonicated for 15 min and centrifuged at 12,000 g for 10 min at 4 °C.

### Metabolomics data acquisition and processing

A 1 µL aliquot of the extract was subjected to LC-MS analysis using the Sciex TripleTOF 6600 system coupled with the Agilent 1290 HPLC. For polar metabolite analysis, mobile phase A was water with 10 mM ammonium formate and mobile phase B was acetonitrile with 0.1% formic acid. Flow rate was 0.250 ml/min. The LC gradient started at 3% A, increased to 30% A from 1 to 12 min, then to 90% A from 12 to 15 min, remained at 90% until 18.5 min, and returned to 3%, holding until the end of the 24-min run. For lipid analysis, mobile phase A was prepared by 60:40 water: acetonitrile with 10 mM ammonium formate, and mobile phase B was made by 90:10 isopropanol: acetonitrile with 10 mM ammonium formate. Flow rate was 0.250 ml/min and the total run time is 15.80 min. The LC gradient started at 80% A, decreased to 40% A at 2 min, further decreased to 0% from 2 to 12 min, remained at 0% A until 14 min, then returned to 80% A at 14.10 min, and held at 80% A until the end of run. Mass spectrometry settings were as follows: gas1 50 V, gas2 60V, curtain gas 25V, source temperature 500 °C, IonSpray voltage 5500 V. The collision energy was set to 30 V and 45 V for polar metabolites and lipids, respectively. Fifteen precursors were selected for MSMS fragmentation in the DDA acquisition. The mass range for the RP method was 300 – 1000 m/z and for the HILIC method was 50 – 1000 m/z. In the SWATH acquisition, fixed windows were applied with window widths of 21 Da for the HILIC method and 10 Da for the RP method.

Each tissue type (i.e., plasma and liver) was analyzed as a separate batch. Following sample reconstitution, the supernatants were collected into MS vials and pooled as quality control (QC) samples for each tissue type. For each analytical batch, patient samples were injected in a randomized sequence, along with one QC injection approximately every 10 samples for monitoring instrument stability and calculating the coefficient of variation (CoV). Both reverse phase (RP) and hydrophilic interaction liquid chromatography (HILIC) columns were used for chromatographic separations of organic and aqueous fractions, respectively. For the analysis of aqueous fraction, metabolites were separated on SeQuant ZIC-cHILIC (3 μm, 100Å, 100×2.1 PEEK) column with a flow rate of 0.25 mL/min in a 24-minute run. For the organic fraction, metabolites were separated on RRHD Eclipse Plus C18 (2.1 x 50 mm, 1.8 μm).

Data acquisition was performed in both positive and negative ionization modes. Data-dependent acquisition (DDA) was performed on pooled QC samples for compound identification and SWATH-MS analysis was applied to individual samples for relative quantification. The mass range for the RP method was 300 – 1000 m/z and for the HILIC method was 50 – 1000 m/z. Data was converted into mzML format and was then processed by MetaboKit (Narayanaswamy et al., 2020) (https://github.com/MetaboKit). To gain broad metabolite coverage, we curated a data processing pipeline for metabolites identified via spectral matches (MSMS-level, with 915 identifications postprocessing) and those identified solely through mass matches (MS1-level, with 548 identifications postprocessing). For metabolite quantification, the project-specific spectral library generated from DDA analysis was used to extract MS1 and MS2 features from the SWATH data using the MetaboKit DIA module. Compound transitions with a CoV > 30% were excluded from the analysis. For each compound, the precursor or fragment ion with the lowest CoV was selected as the quantifier. For a compound identified in both positive and negative modes using the same column, the peak feature with the lower CoV were selected. Overlapping metabolites in the RP and HILIC datasets are mainly semi-polar lipids, as RP detects compounds with m/z > 300. For lipids identified by both methods, the ones measured on the RP column was selected.

### RNA sequencing

Transcriptome sequencing service was provided by Biomarker Technologies (BMK), GmbH. Briefly, RNA were extracted from liver biopsies using Trizol reagent. The quality and quantity of extracted RNA was assessed by Nanodrop (quantity and purity) and Labchip GX (Quality). The cDNA libraries were prepared using Hieff NGS Ultima Dual-mode mRNA Library Prep Kit from Illumina (Yeasen) as per manufacturer’s instructions. RNA sequencing was performed on NoveSeq6000 platform (PE 150 mode). For raw data processing, FastQC (v0.11.9) was used as a quality control check for raw sequencing. Alignment to the reference genome (GRCh38, Ensembl release 77) was performed using STAR (v2.7.9a), and gene expression levels (transcripts-per-million, TPM) were estimated using RSEM (v1.3.1). Gene expression data underwent a logarithmic transformation to log2(TPM + 1) and mean-centering normalization for downstream analyses.

### Statistical analysis

All analyses in this study were conducted using R version 4.3.1. Linear regression model was used to evaluate the linear association between histological grades and molecular abundance levels with adjustment for age, sex, BMI, and diabetes. Statistically significantly changed metabolites or genes were defined by controlling the overall type I error at 5% or 10% (q value < 0.05 or 0.1). Spearman rank correlation was used to assess the relationship between the same metabolite in the liver and plasma. This accounts for matrix effects across different tissue types in untargeted analysis by comparing the relative intensity ranking of a metabolite across all samples within each tissue type. Gene overrepresentation analyses were performed using gene ontology recourse by in-house software (Gene Ontology et al., 2023). ClueGO, a plugin app in Cytoscape, was used to visualize enrichment maps (Bindea et al., 2009). Partial correlation network analysis was employed to integrate the two omics layers using an in-house partial correlation network algorithm ACCORD (Lee et al., 2025). KEGG pathway database was used for the analysis of lipid metabolism and pathway mapping (Kanehisa et al., 2016). All network visualizations in this study were done using Cytoscape (Otasek et al., 2019).

To investigate MASLD subgroup signatures related to diabetic status, we analyzed linear associations between gene signatures and histological features separately in non-diabetic (n = 71) and diabetic individuals (n = 23). Statistical power was estimated by comparing variance explained in full (y ∼ x + a + b + c) versus reduced (y ∼ a + b + c) regression models, converting the incremental *R^2^* into Cohen’s *f^2^*, and applying pwr.f2.test at α = 0.05. We further compared gene expression profiles between diabetic (n = 21) and non-diabetic (n = 43) MASLD patients, identifying 166 differentially expressed genes (p < 0.05, |log₂FC| > 0.32). Of these, 54 genes (**Supplementary file 10**) were both differentially expressed and significantly associated with fibrosis progression, and thus marked as signatures in diabetic MASLD.

To identify progressive markers of liver fibrosis, we performed pairwise analyses for fibrosis stages using two distinct baselines: patients with no MASLD and those with steatosis but no fibrosis. 213 markers were characterized as progressive based on the following criteria: a) significant linear association with fibrosis grades, and b) significant differential expression (p value < 0.05 in t-test) in at least two fibrosis stages (e.g., F1 and F2, F2 and F3) compared to both baselines.

To identify independent gene signatures associated specifically with steatosis and fibrosis progression, we applied linear regression models for each outcome adjusting for the other. Steatosis-specific signatures were defined as genes significantly associated with steatosis but not with fibrosis, and vice versa. Metascape was used for meta-analysis of enrichment pathways (Zhou et al., 2019). Pathways with at least 10 hits and p value < 0.05 were considered steatosis- or fibrosis-specific.

For meta-analysis of reference human cohorts, the two RNA-seq datasets of the liver transcriptome, the VA cohort (GSE130970) and the EU cohort (GSE135251), were downloaded from Gene Expression Omnibus, and associations between gene expression (log2 TPM) and fibrosis grades or NAS were analyzed using linear regression. In the EU cohort, the patients without MASH diagnosis were removed from the meta-analysis for the purpose of identifying fibrosis signatures in MASH patients. Human liver single cell dataset was obtained from GSE115469 (MacParland et al., 2018). The mean log2 TPM was calculated for each cell type, and cell specificity was determined by subtracting the mean expression levels across all cell types.

### Cross-referencing datasets of mouse models and human liver organoids

The RNA sequencing data (raw counts) for HSCs isolated from liver fibrosis mouse models (De Smet et al., 2021) was downloaded from GSE176042 and analyzed using the DESeq2 R package. This previously published study investigated HSC initiation and perpetuation using acute and chronic models, respectively. For the acute model, mice were injected with CCl_4_ once and samples were taken at 24 h, 72 h, and 1 w. For the chronic model, mice received a regimen of semi-weekly injection of CCl_4_ for 4 weeks. After this regimen, mice were sacrificed and samples were taken at 24 h, 72 h, and 2 w. The statistical results from single cell dataset of stem cell-derived human liver organoids were obtained from GSE207889 (Hess et al., 2023).

### 3D liver spheroid MASH model

3D liver spheroids were generated based on co-cultures of cryopreserved primary human hepatocytes (PHH) and liver non-parenchymal cells (NPC) as previously described (Youhanna et al., 2025). Briefly, cells were seeded at a PHH:NPC ratio of 6:1 in 96-well ultra-low attachment plates (CLS7007-24EA Corning) with a total of 1500 cells/well in culture medium (William’s E medium containing 11 mM glucose, 100 nM dexamethasone, 10 μg/mL insulin, 5.5 mg/L transferrin, 6.7 μg/L selenite, 2 mM L-glutamine, 100 U/mL penicillin, 0.1 mg/mL streptomycin) supplemented with 10% FBS. After spheroid formation FBS was phased out and cells were maintained in medium supplemented with albumin-conjugated free-fatty acids (FFA) including 80μM palmitic acid and 80 μM oleic acid. At the end of the experiment, spheroids were collected for RNA extraction (Zymo R105). The elafibranor treatment protocol was described in the previous study (Youhanna et al., 2025). Each experimental condition (control, MASH, and elafibranor treatment) was performed in quadruplicate. The functional characterization of hepatocytes in the liver spheroid culture model has been published previously (Bell et al., 2018; Bell et al., 2016; Bell et al., 2017; Messner et al., 2018; Vorrink et al., 2017).

## Acknowledgments

This manuscript is the result of a wonderful international collaboration and we thank all present and past members of the Watt, Choi, Lauschke, and Kaldis laboratories for discussions, input, and support. P.K. thanks Åke Nilsson for discussions and suggestions.

## Abbreviations

ALT: Alanine aminotransferase
AST: Aspartate aminotransferase
ALP: Alkaline phosphatase
CAR: Acylcarnitines
CCl_4_: Carbon tetrachloride
Cer: Ceramide
CM: Conditioned medium
CYP: Cytochrome P
DAG/DG: Diacylglycerol
DDA: Data-dependent acquisition
ECM: Extracellular matrix
FBS: Fetal bovine serum
FFA: Free-fatty acids
FXR: Farnesoid X receptor
GAP: GTPase-activating protein
GEF: Guanine nucleotide exchange factor
GGT: Gamma-glutamyltransferase
GL: Glycerolipid
GPL: Glycerophospholipid
HSC: Hepatic stellate cell
HILIC: Hydrophilic interaction liquid chromatography
LD: Lipid droplet
LNAPE: N-acyl-lysophosphatidylethanolamine
LPC-O: Ether-linked lysophosphatidylcholine
LPI: Ether-linked lysophosphatidylinositol
LPS: Ether-linked lysophosphatidylserine
LPC: Lysophosphatidylcholine
LPE: Lysophosphatidylethanolamine
MG: Monoacylglycerol
MASH: Metabolic dysfunction-associated steatohepatitis
MASLD: Metabolic dysfunction-associated steatotic liver disease
NAS: NAFLD activity score
NPC: Non-parenchymal cells
NAE: N-acyl ethanolamines
PCA: Principal component analysis
PHH: Primary human hepatocytes
PUFA: Polyunsaturated fatty acid
PC: Phosphatidylcholine
PC-O: Ether-linked phosphatidylcholine
PE: Phosphatidylethanolamine
PE-O: Ether-linked phosphatidylethanolamine
PG: Phosphatidylglycerol
PI: Phosphatidylinositol
PS: Phosphatidylserine
QC: Quality control
RP: Reverse phase
SM: Sphingomyelin
SL: Sphingolipid
SPB: Sphingoid base
SAM: S-adenosylmethionine
SAH: S-adenosylhomocysteine
TAG/TG: Triacylglycerol/Triglyceride
TAG-O: Ether-linked triacylglycerol
Vitamin E: Alpha-tocopherol

## Additional files

Supplementary files

Supplementary file 1. Additional patient characteristics.

Supplementary file 2. Top-enriched pathways of genes with nonzero loading scores on PC1 and PC2 in the sparse PCA of the liver transcriptome.

Supplementary file 3. Statistical analysis of metabolomics data for plasma samples.

Supplementary file 4. Statistical analysis of metabolomics data for liver samples.

Supplementary file 5. Statistical analysis of transcriptomics data for liver samples

Supplementary file 6. Associations between mitochondrial function-related genes and liver histological grades.

Supplementary file 7. Associations between autophagy-related genes and liver histological grades.

Supplementary file 8. Over-representation analysis enrichment analysis of genes linearly associated with steatosis or fibrosis grades.

Supplementary file 9. Steatosis-specific and fibrosis-specific gene signatures

Supplementary file 10. Subgroup statistical analysis of liver transcriptome in diabetic and non-diabetic individuals.

Supplementary file 11. Progressive gene markers associated with liver fibrosis in MASLD patients.

Supplementary file 12. Genes involved in the transition from fibrosis-free steatosis to fibrosis.

Supplementary file 13. Associations between GTPase-related genes and liver histological grades.

Appendix 1 – supplementary methods

Appendix 2 – discussion of plasma metabolome

## Data availability

All data are available within the manuscript, supplementary figures, and supplementary tables. RNA sequencing data to this article have been submitted to SRA (PRJNA1185558) and is deposited with GEO (GSE281797). Untargeted LC-HRMS data is deposited with Zenodo, including IDA data (DOI 10.5281/zenodo.14091962), SWATH data for the organic fraction (DOI 10.5281/zenodo.14096635), and SWATH data for the aqueous fraction (DOI 10.5281/zenodo.14096753, DOI 10.5281/zenodo.14136832). All clinical data, processed omics datasets, and code are available at https://github.com/SLINGhub/MASLD_dual_omics.

The following datasets were generated:

| Author(s) | Year | Dataset title | Dataset URL | Dataset and identifier |
| --- | --- | --- | --- | --- |
| Zhao Q, De Nardo W, Choi H, Watt MJ, Kaldis P | 2025 | Transcriptomic profiles of liver biopsies in obese patients with metabolic dysfunction-associated steatotic liver disease | https://www.ncbi.nlm.nih.gov/geo/query/acc.cgi?acc=GSE281797 | NCBI Gene Expression Omnibus, GSE281797 |
| Zhao Q, De Nardo W, Choi H, Watt MJ, Kaldis P | 2024 | Untargeted LC-HRMS-based metabolomic profiling reveals distinct metabolomic profiles in patients with metabolic dysfunction-associated steatotic liver disease (MASLD) | https://zenodo.org/records/14091963 | Zenodo DOI 10.5281/zenodo.14091962, DOI 10.5281/zenodo.14096635, DOI 10.5281/zenodo.14096753, DOI 10.5281/zenodo.14136832 |

## Author contributions

QZ*: designing research studies, conducting experiments, acquiring data, analyzing data, writing the manuscript

WD*: designing research studies, conducting experiments, acquiring data, analyzing data, writing the manuscript

RW*: analyzing data, writing the manuscript

YZ: conducting experiments, acquiring data, analyzing data

UK: analyzing data, writing the manuscript

GS: conducting experiments, acquiring data, analyzing data

LNZ: analyzing data, writing the manuscript

HT: conducting experiments, acquiring data, analyzing data

SY: conducting experiments, acquiring data, analyzing data

MY: conducting experiments, acquiring data, analyzing data

YX: conducting experiments, acquiring data, analyzing data

YK: conducting experiments, acquiring data, analyzing data

SL: conducting experiments, acquiring data, analyzing data

RL: conducting experiments, acquiring data, analyzing data

GT: conducting experiments, acquiring data, analyzing data

PN: conducting experiments, acquiring data, analyzing data

PRB: providing reagents, cohort management

VMK: designing research studies, analyzing data, writing the manuscript, supervision, funding acquisition

HC: designing research studies, analyzing data, writing the manuscript, supervision, funding acquisition

MJW: designing research studies, analyzing data, writing the manuscript, supervision, funding acquisition

PK: designing research studies, analyzing data, writing the manuscript, supervision, funding acquisition, leading of collaboration

QZ, WD, and RW are co-first authors based on their contributions, which was writing original draft, LC-MS data acquisition and bioinformatic analysis of transcriptomic, metabolomic, and clinical data (QZ), organizing cohort samples, clinical parameters, managing, and LX-2 experiments (WD), and writing the original draft and analyzing data (RW).

---

**Figure 2–figure supplement 1.**
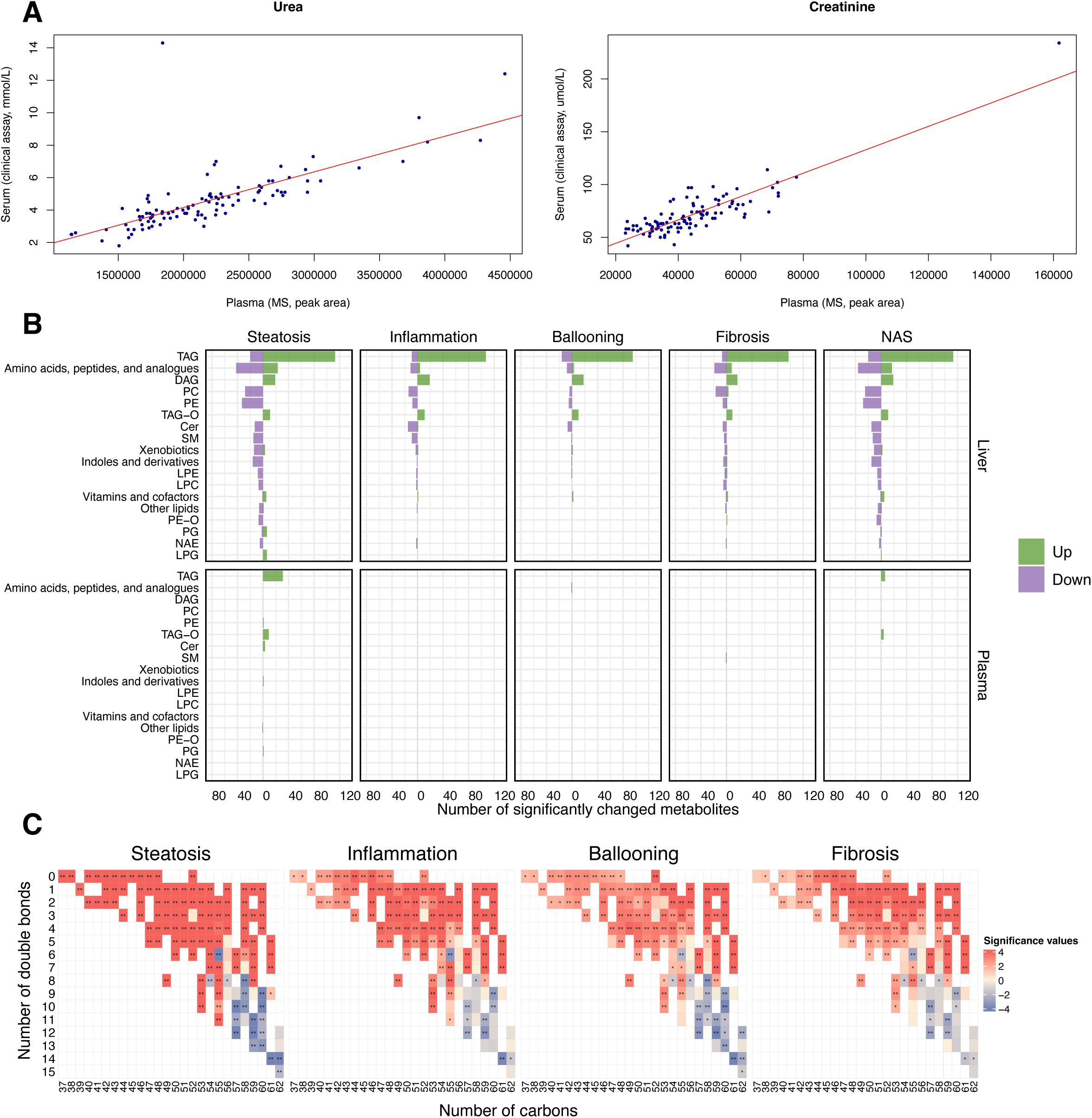
Metabolomic analysis in this obese MASLD cohort. (A) Plasma urea and creatinine from untargeted metabolomics correlated with clinical assays. (B) Metabolites associated with the disease progression in each matrix (q < 0.05 or q < 0.1). Compound classes with at least 5 hits associated with any histological feature were shown in the plot. (C) Associations between fatty acid composition of triacylglycerides (TAGs) and histological outcomes. Significance values refer to –log10(p value)*sign(coefficient) from linear regression models.

**Figure 2–figure supplement 2.**
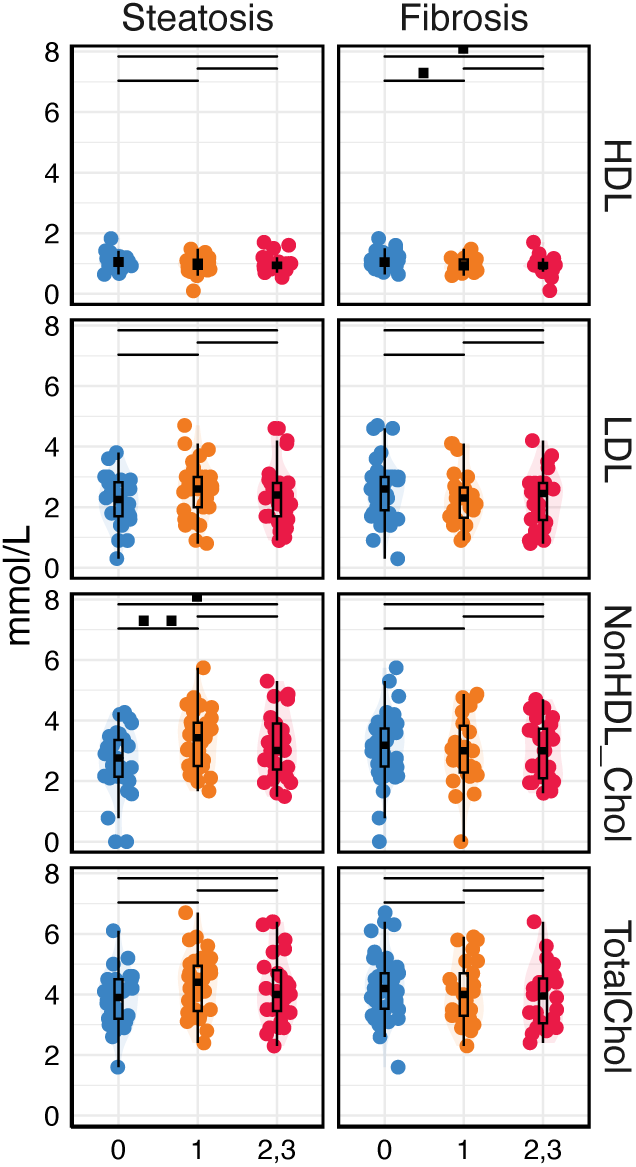
Blood lipoprotein cholesterol levels in patients with different steatosis and fibrosis grades. HDL, high-density lipoprotein cholesterol. LDL, low-density lipoprotein cholesterol. NonHDL-Chol, non-high-density lipoprotein cholesterol. TotalChol, total blood lipoprotein cholesterol.

**Figure 3–figure supplement 1.**
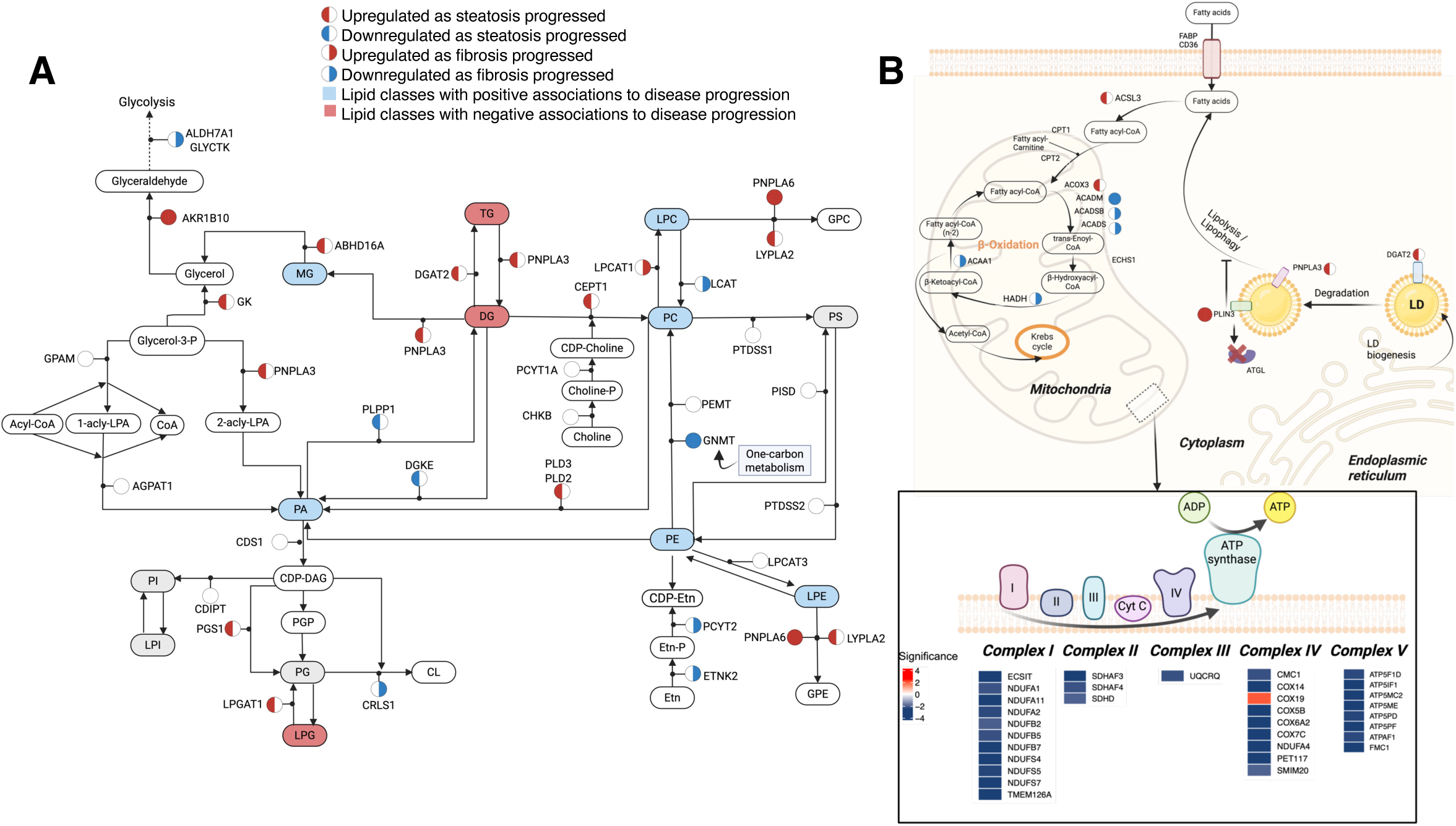
Integrative map of liver metabolism in obese individuals with MASLD. (A) An integrative map of transcriptomics and metabolomics of glycerolipid and glycerophospholipid metabolism in early MASLD. (B) Gene expression changes in fatty acid β-oxidation, mitochondrial respiratory chain, and lipid droplet (LD) metabolism associated with steatosis and fibrosis. There was a notable downregulation of genes involved in the electron transport chain of the inner mitochondrial membrane, specifically in complex I (NDUF family), complex II (SDH family), complex III (UQCR), complex IV (COX), and complex V (ATP5 family) as fibrosis progressed. Genes involved in respiratory electron transport were colored by the significance (–log10(p value)*sign(coefficient)) in relation to fibrosis grades. Inset: gene expression changes in the electron transport chain are depicted.

**Figure 3–figure supplement 2.**
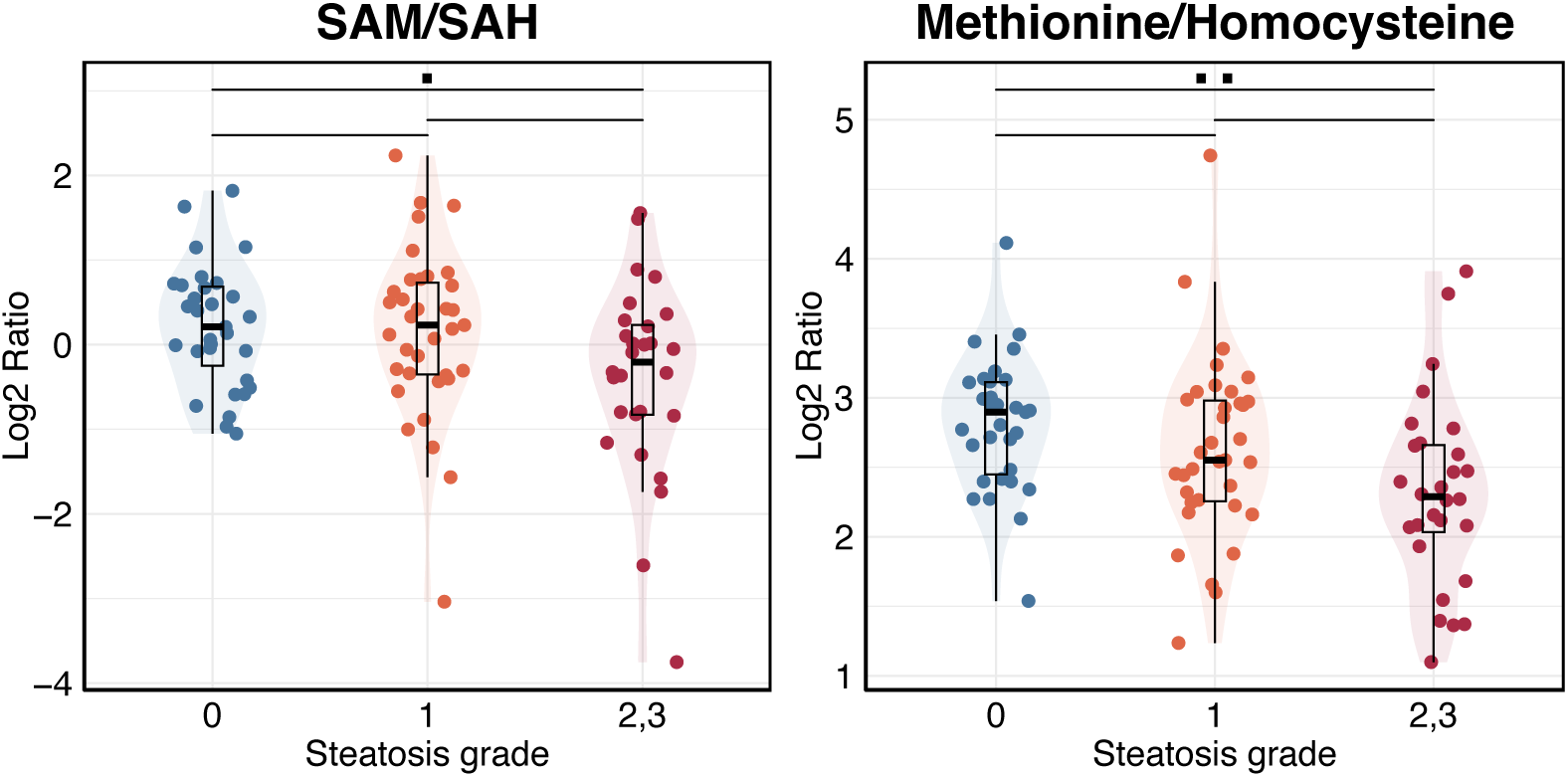
Ratios of metabolites in one-carbon metabolism in individuals with different steatosis grades. SAM, S-adenosylmethionine. SAH, S-adenosylhomocysteine.

**Figure 4–figure supplement 1.**
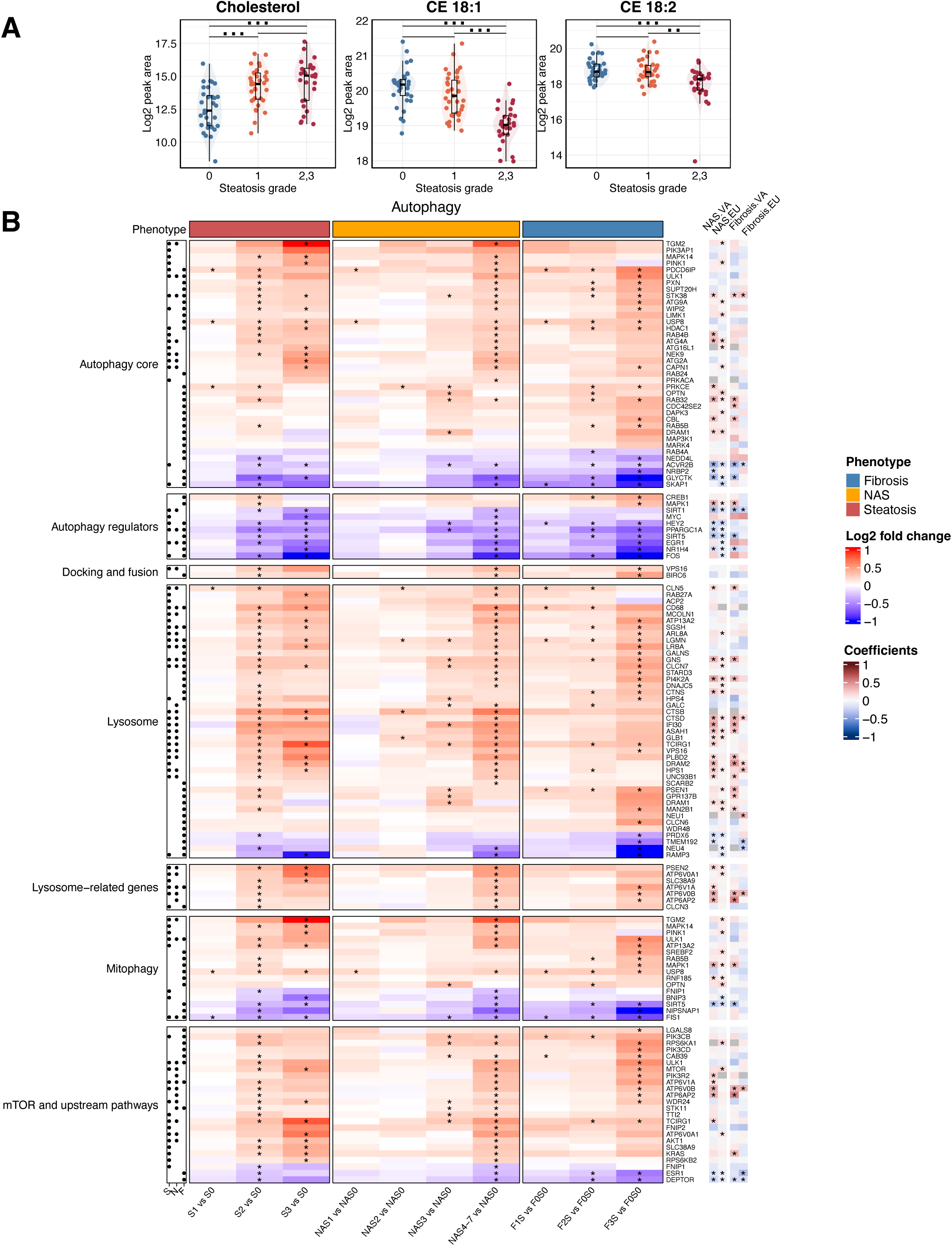
Autophagy regulation in the liver. (A) Hepatic free cholesterol, CE 18:1, and CE 18:2 levels in this cohort in relation to steatosis grade (x-axis). (B) Heatmap of autophagy genes in relation to histological features, indicating active lipophagy in the liver of obese individuals. Asterisks in the main heatmap denote the significance (p < 0.05) from comparisons of each subgroup. Association with steatosis (S), NAS (N) and fibrosis (F) was indicated by black dots (q < 0.05, left). Results were cross-referenced with two published cohorts (VA cohort, GSE130970; EU cohort, GSE135251) (*q < 0.05 and with consistent direction of changes as in our cohort, bottom).

**Figure 5–figure supplement 1.**
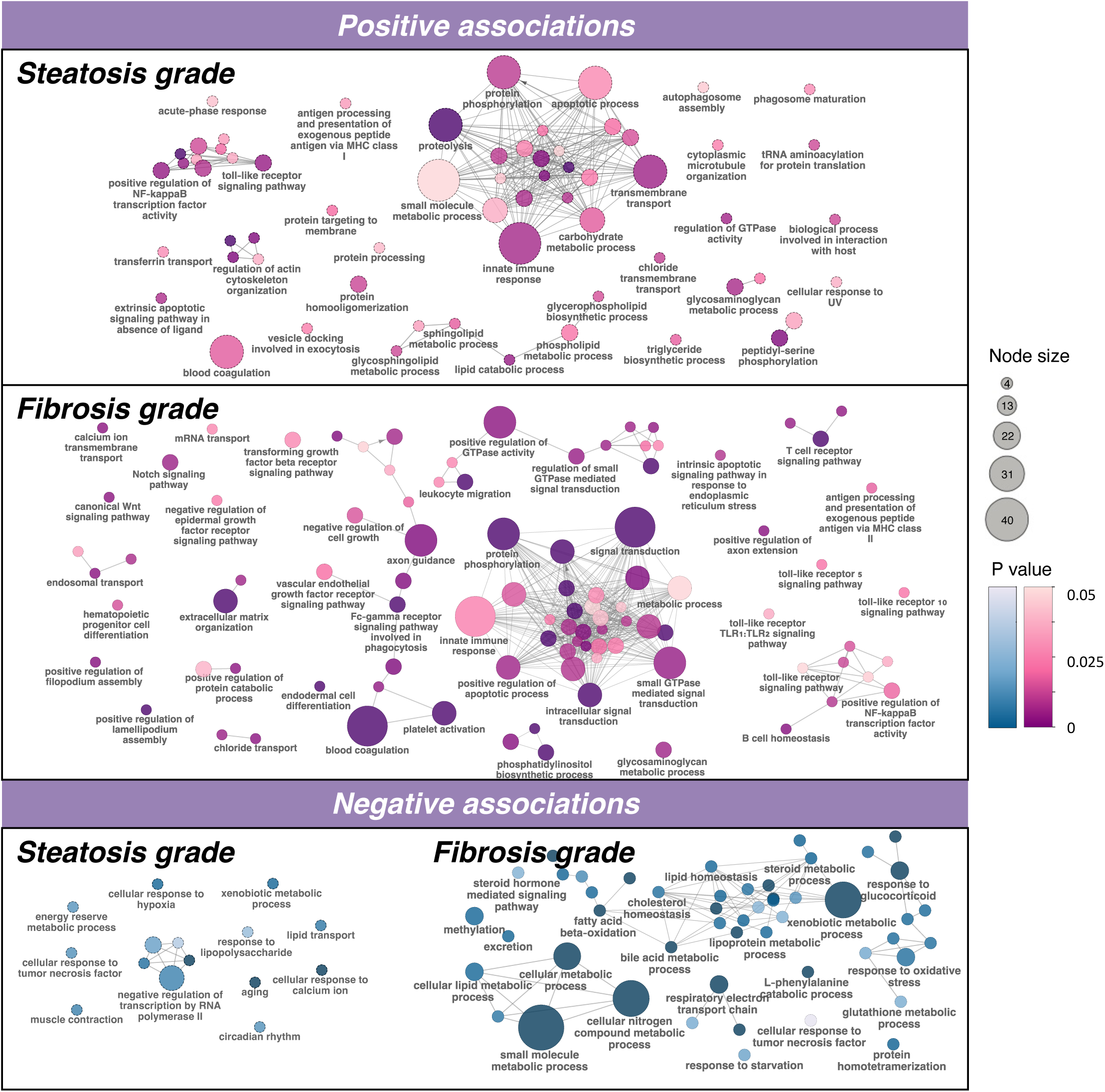
Functional enrichment of gene sets associated with steatosis and fibrosis in the liver transcriptome. Node size is depicted by circles and P value by color.

**Figure 5–figure supplement 2.**
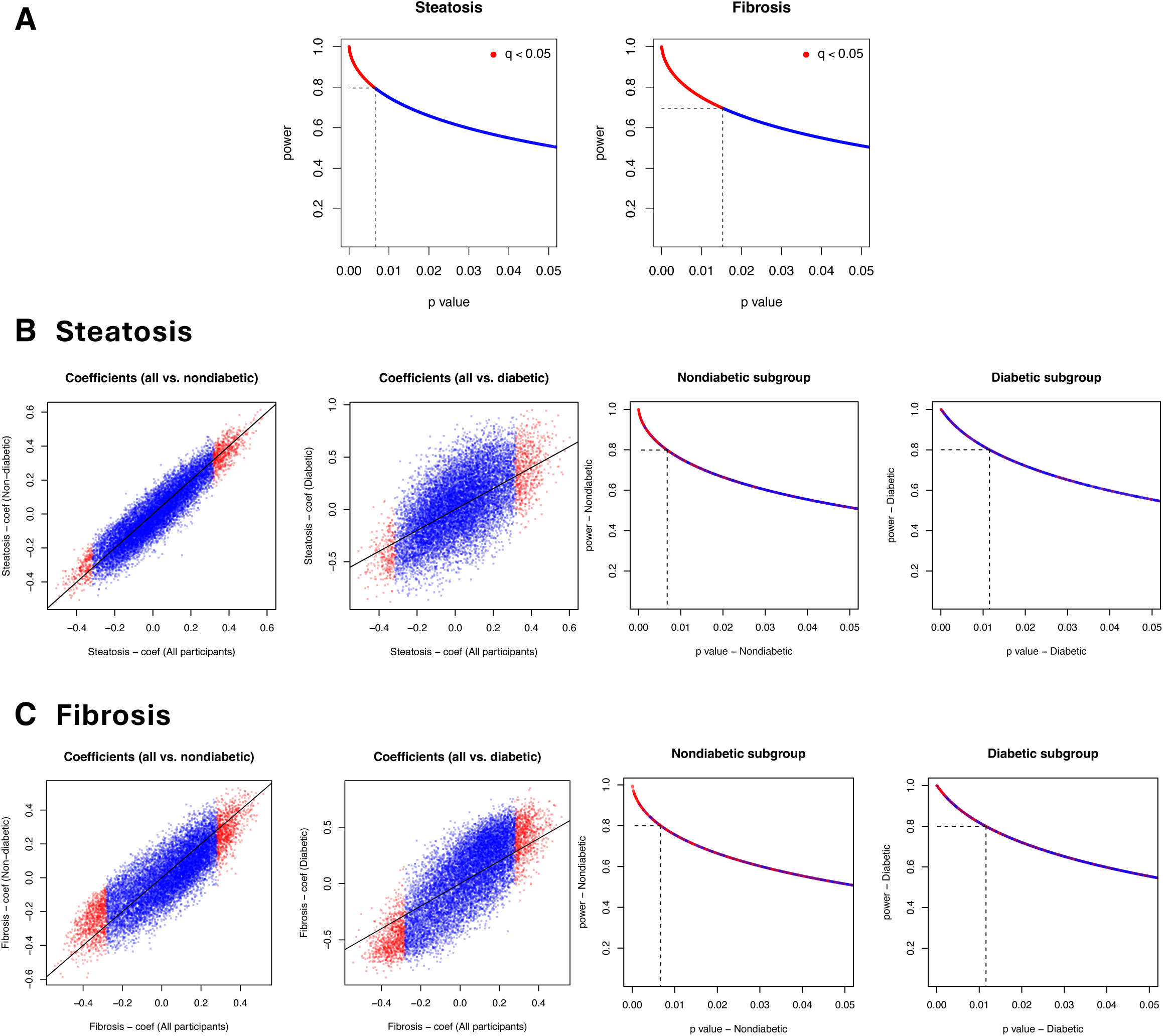
Statistical power and subgroup analysis of associations between gene expressions and steatosis or fibrosis grades. Statistical power in the original linear regression analysis of steatosis and fibrosis was plotted against p-values (A). Differential features associated with steatosis (q < 0.05) showed power > 0.8, whereas those associated with fibrosis showed power > 0.7. Subgroup analyses of participants with and without diabetes were conducted to evaluate associations between gene expression and steatosis (B) and fibrosis (C). The x-axis shows coefficients from the original analysis including all participants. Corresponding coefficients from the analyses of non-diabetic (left panel) and diabetic (middle-left panel) individuals are displayed alongside the statistical power for each subgroup analysis (middle-right and right panels). Red dots represent genes that were significant in the original analysis (q < 0.05). Subgroup analyses suggest that the results in the non-diabetic subgroup (n = 71) were highly consistent with findings from the original analysis (n = 94, adjusted for diabetes), indicating that the originally reported gene signatures, after correction for diabetic status, remain valid in non-diabetic individuals.

**Figure 6–figure supplement 1.**
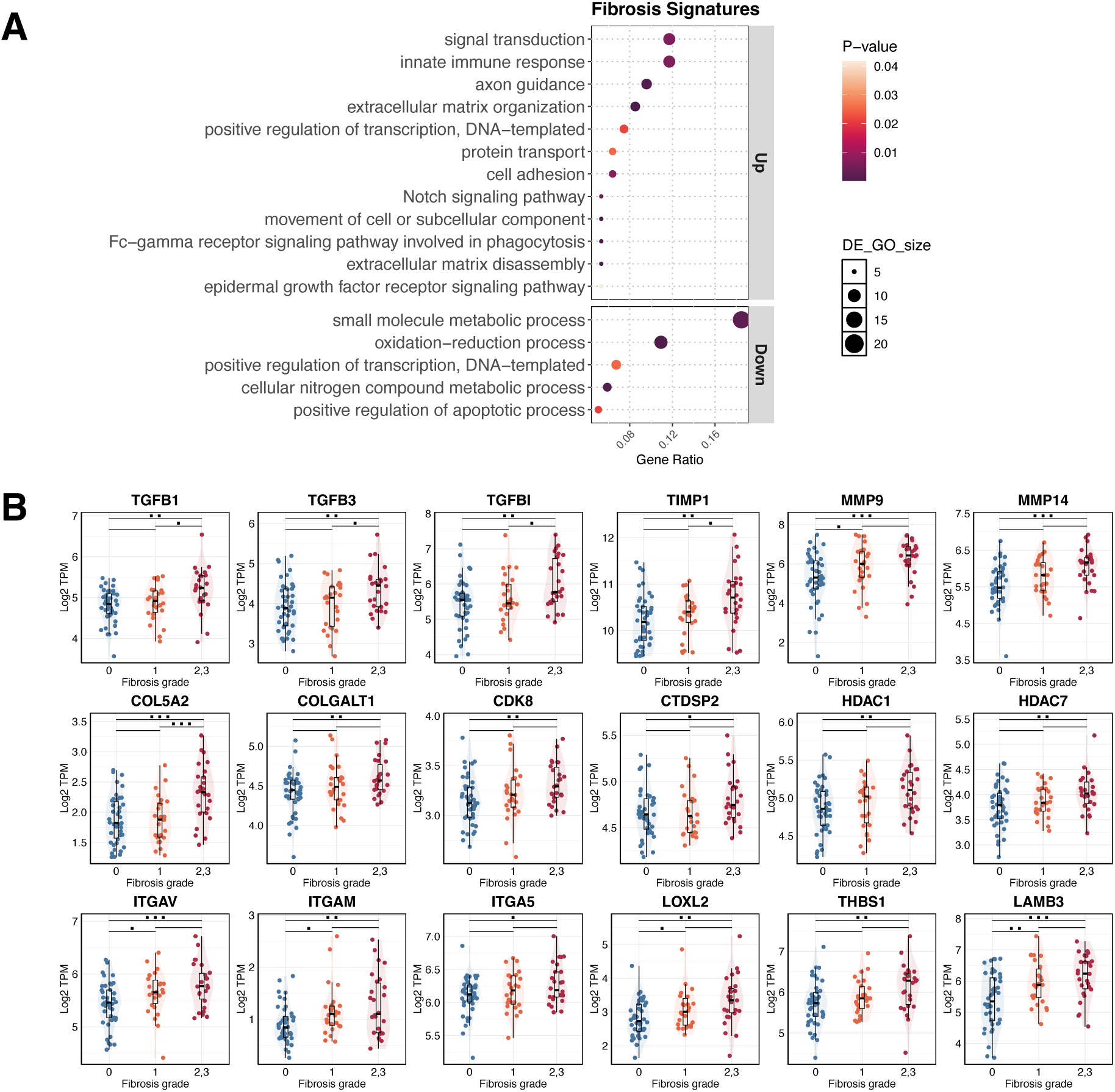
Liver fibrosis pathways and gene signatures. (A) Pathway analysis of the enrichment of pathways for the 213 fibrosis signatures. (B) TGF-β and SMAD related gene expression associated with fibrosis grades (x-axis) is depicted. Significance is indicated by black dots; * p < 0.05, ** p < 0.01, *** p < 0.001.

**Figure 6–figure supplement 2.**
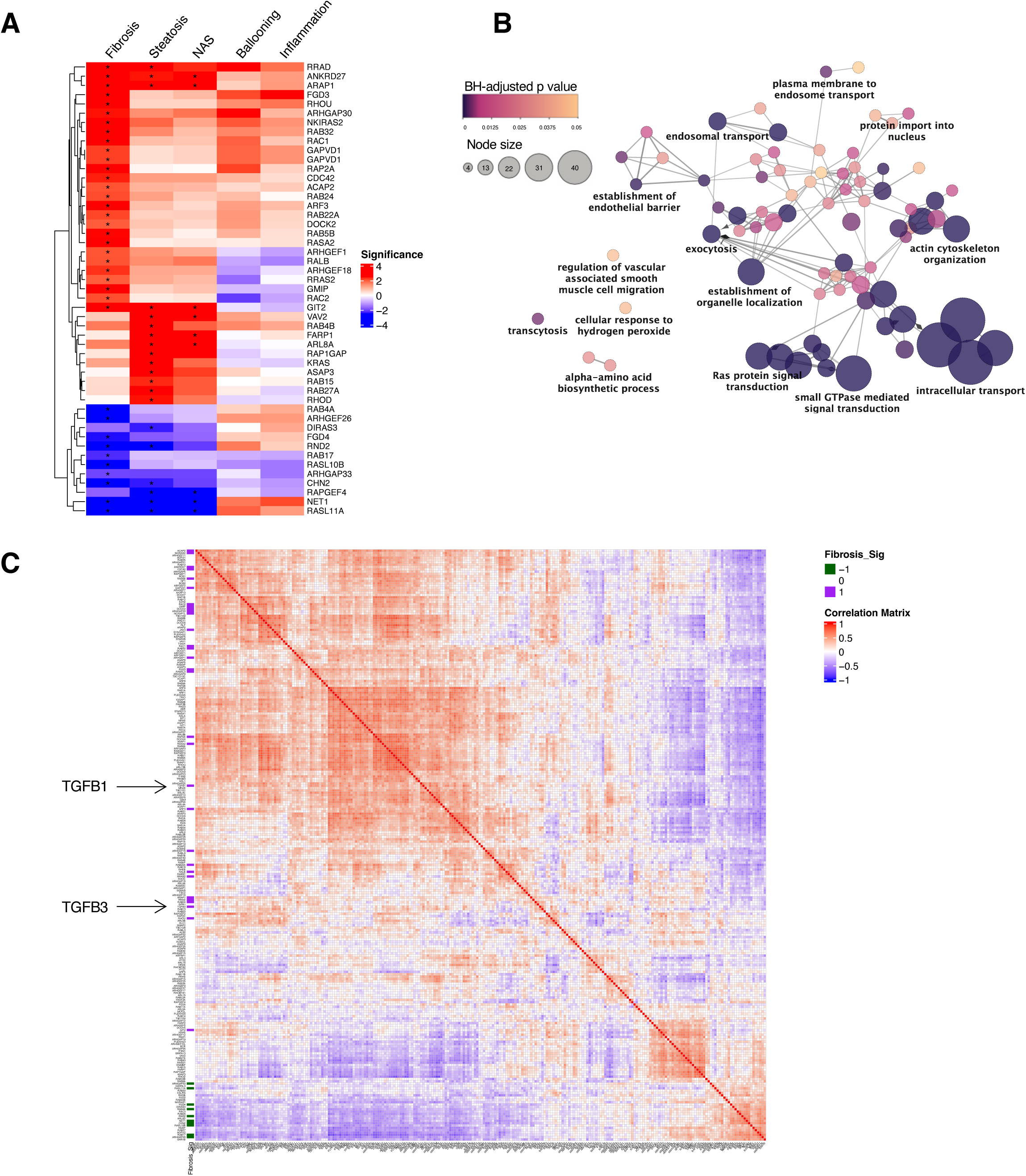
GTPases and their regulation emerges as a potential target for liver fibrosis. (A) Heatmap of GTPase-related genes display a better correlation with fibrosis grades. Asterisks in the main heatmap denote the significance (p < 0.05) from comparisons of each subgroup. (B) Enrichment map of selected genes constituting the core of the protein-protein interaction network of GTPase-related genes. Nodes are colored according to BH (Benjamini-Hochberg) adjusted p-value, and node sizes are proportional to the number of genes in the term. (C) Correlation matrix of gene expression levels for TGF-β and GTPase-related genes. The correlation plot was created of Pearson’s r values for GTPase-related genes and TGF-β genes (TGFB1 and TGFB3). The left annotation bar indicates the significance of associations between g7ene expression and fibrosis grades.

**Figure 6–figure supplement 3.**
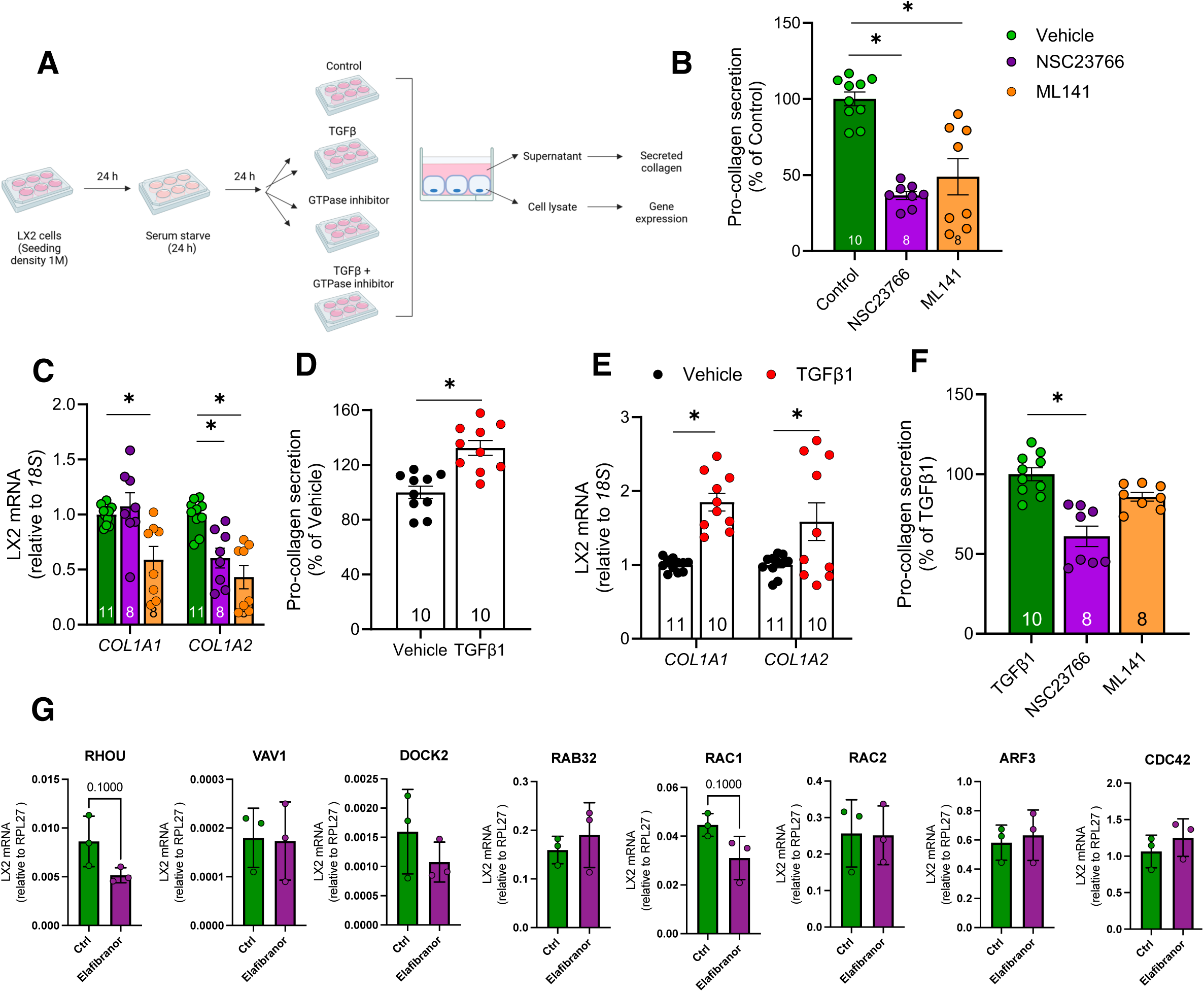
GTPase inhibition and Elafibranor treatment in LX2 cells. (A) Workflow of the LX2 experiment in panel A-F (n = 8-10). (B) GTPase inhibitors NSC23766 (Rac1 inhibitor) and ML141 (Cdc42 inhibitor) significantly reduced pro-collagen secretion from HSC-like LX-2 cells and (C) inhibited gene expression of COL1A1 and COL1A2 under basal conditions. (D) TGFβ administration increased pro-collagen secretion by 32% and (E) collagen gene expression in LX2 cells. (F) NSC23766-mediated GTPase inhibition impairs pro-collagen secretion from LX-2 cells after TGF-β treatment. Pro-collagen secretion was determined by ELISA and gene expression levels were assessed by qPCR. Asterisks (*) denote p value < 0.05 for statistical significance from one-way ANOVA with Holms-Sidak multiple comparisons (B&C), unpaired t-test (D&E) Kruskal Wallis Test with Dunns multiple comparisons (F). (G) Gene signatures in LX-2 cells with and without Elafibranor (n = 3).

**Figure 6–figure supplement 4.**
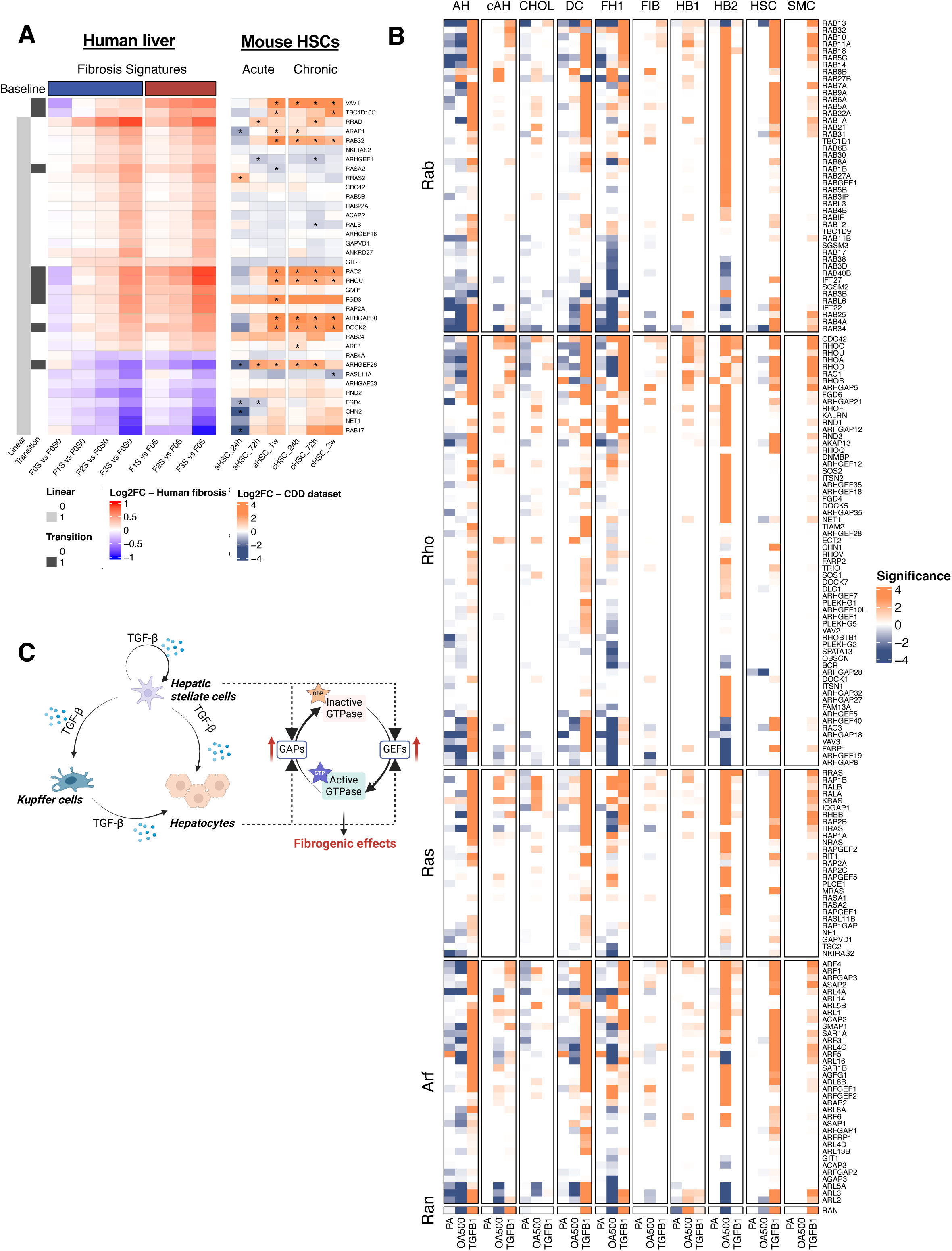
GTPase-related genes in external systems. (A) Human fibrosis signatures from the present study were partially reflected in a mouse study (GSE176042), where RNA sequencing was performed on hepatic stellate cells (HSCs) isolated from both acute and chronic mouse fibrosis models. GTPase-related genes, especially those involved in the initiation of fibrosis, were also upregulated in HSCs isolated from both acute and chronic liver fibrosis mouse models at different time points. Row annotations ‘Linear’ and ‘Transition’ indicate whether gene expression is linearly associated with fibrosis grades (Linear = 1) or shows a significant difference between patients with fibrosis and steatosis versus simple steatosis (Transition = 1). Asterisk denotes genes with adjusted p value < 0.05. (B) GTPase-related genes induced by TGF-β were elevated in hepatocytes and HSCs at different stages of a human liver organoid system (GSE207889). Significance was calculated by –log10(p value)*sign(fold change). Asterisk denotes genes with adjusted p value < 0.05. AH, adult hepatocyte-like; CHOL, cholangiocyte-like; DC, ductal cell-like; FH1, fetal hepatocyte 1-like; FIB, fibroblast-like; HB1, hepatoblast 1-like; HB2, hepatoblast 2-like; HSC, hepatic stellate cell-like; SMC, smooth muscle cell-like. (C) Schematic illustration of the hypothesized role of GTPase signaling in regulating liver fibrogenesis. TGF-β production is initiated in non-parenchymal liver cells, leading to the overexpression of GTPase regulators in both hepatic stellate cells and hepatocytes. This activation of GTPase signaling amplifies downstream fibrogenic effects, creating a feed-forward loop that intensifies fibrosis progression. Further investigation is needed to prove by which mechanisms GTPases regulate collagen expression and therefore fibrosis.

**Figure 6–figure supplement 5.**
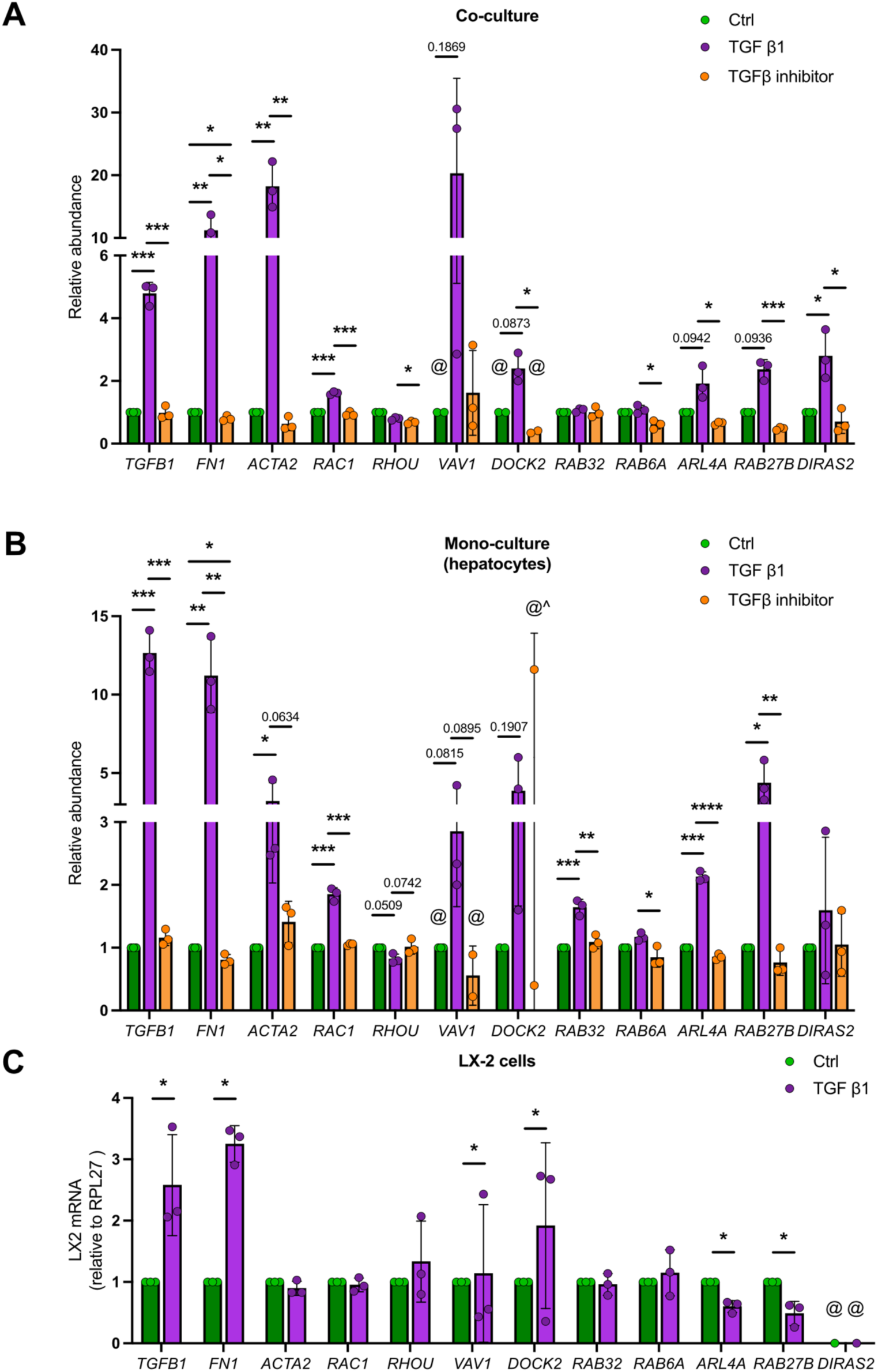
Expression of GTPase-related genes in spheroid co-culture, hepatocyte monoculture, and LX-2 cells. RNA-sequencing-based expression of GTPase-related genes in spheroid co-culture, hepatocyte monoculture upon control (Ctrl), TGF-β1 or TGFβ – inhibitor treatment (A, B). Expression of GTPase-related genes in control (Ctrl) and TGF-β1- treated (24h) LX2 cells measured by qPCR (C). Asterisks (*) denote p value < 0.05, (**) p < 0.005, (***) p < 0.0001 for statistical significance from unpaired t-test, numerical values above the bars indicate p-value. Bars were removed where data did not show strong directionality and statistical reliability, as indicated by (^) symbol. Symbol (@) indicates one or more expression values were not detected in the original dataset. Please note that ACTA2 expression in LX-2 cells is time dependent (unpublished results) and therefore we also included fibronectin (FN1) as a control for TGF-β1 treatment.

