## Supplementary material for "Global molecular landscape of early MASLD progression in obesity": Supplmental Figures

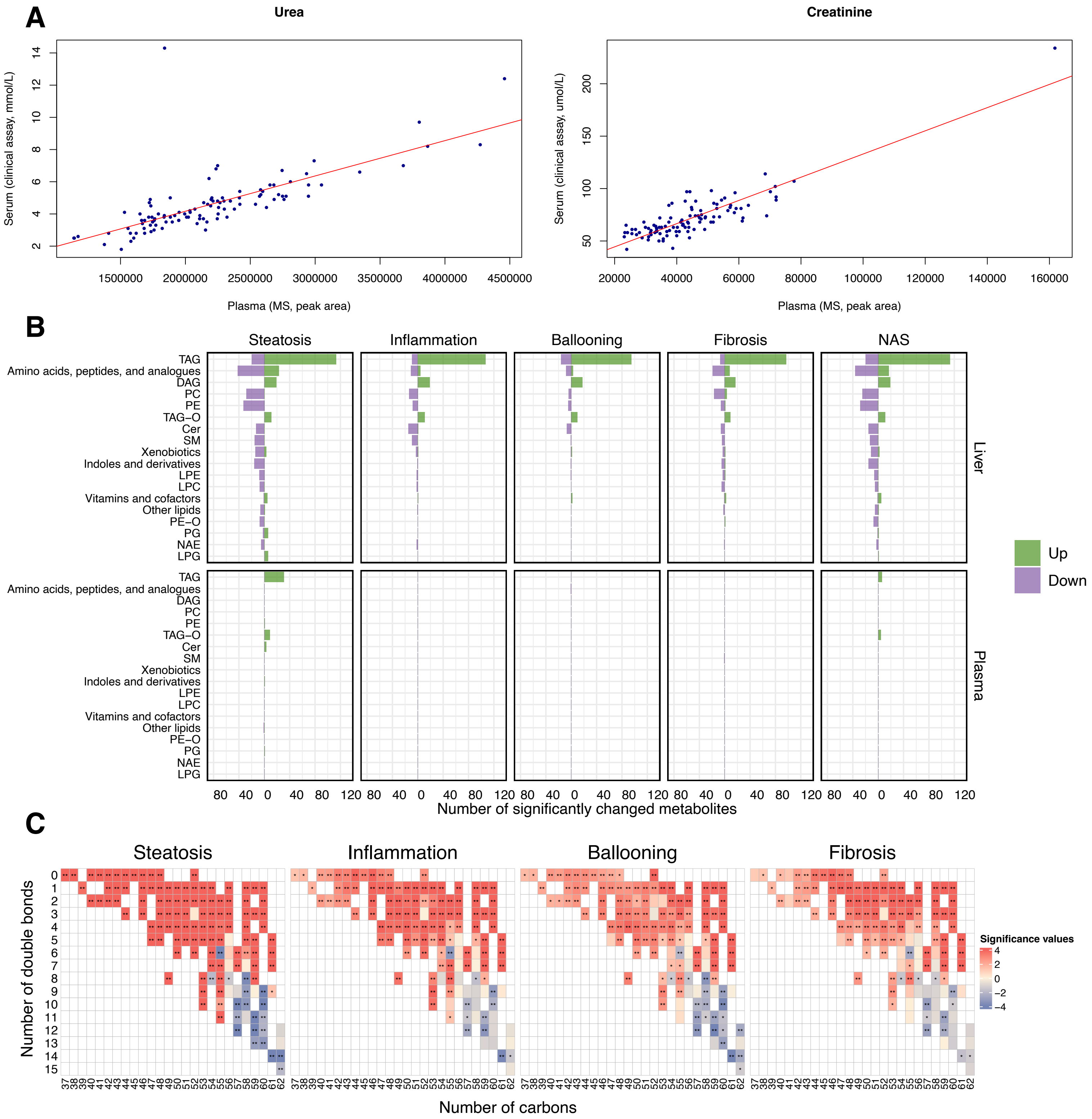


**Figure 2–figure supplement 1. Metabolomic analysis in this obese MASLD cohort**

(A) Plasma urea and creatinine from untargeted metabolomics correlated with clinical assays. (B) Metabolites associated with the disease progression in each matrix ($q < 0.05$ or $q < 0.1$). Compound classes with at least 5 hits associated with any histological feature were shown in the plot. (C) Associations between fatty acid composition of triacylglycerides (TAGs) and histological outcomes. Significance values refer to –log10(p value)*sign(coefficient) from linear regression models.

**Figure 2–figure supplement 2. Blood lipoprotein cholesterol levels in patients with different steatosis and fibrosis grades.** HDL, high-density lipoprotein cholesterol. LDL, low-density lipoprotein cholesterol. NonHDL-Chol, non-high-density lipoprotein cholesterol. TotalChol, total blood lipoprotein cholesterol.

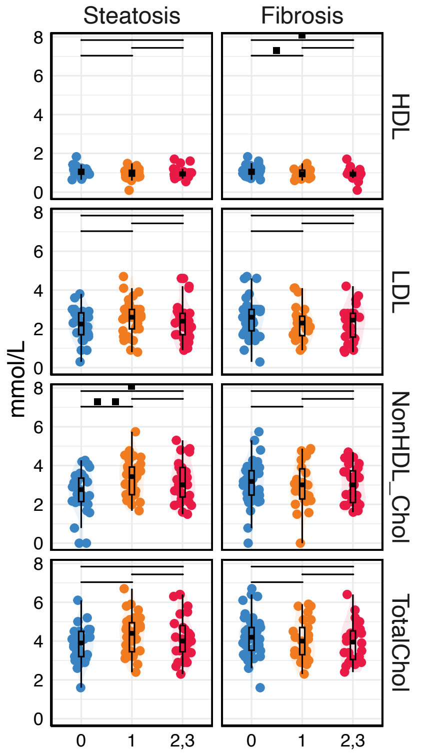


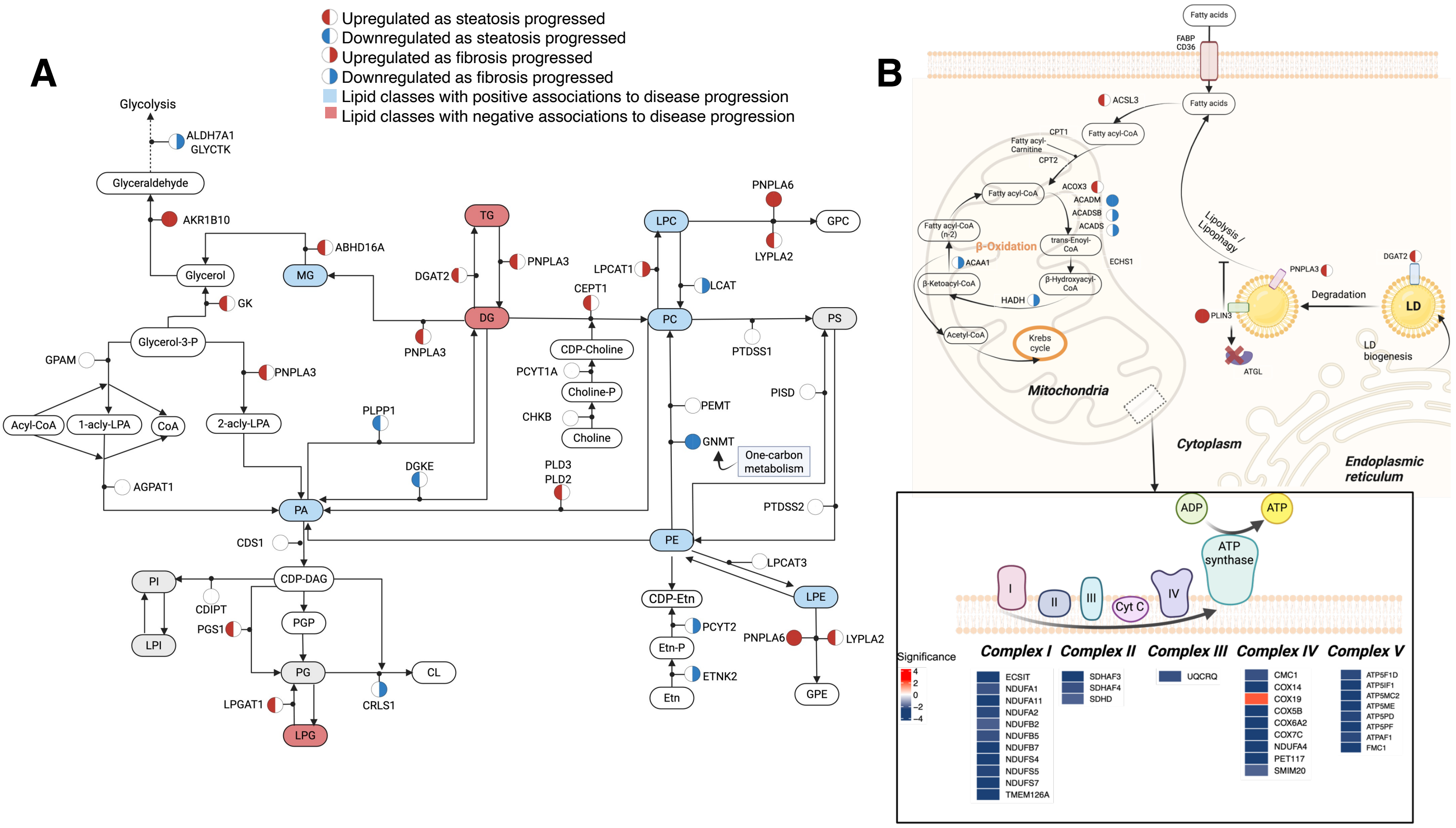


**Figure 3–figure supplement 1. Integrative map of liver metabolism in obese individuals with MASLD.** (A) An integrative map of transcriptomics and metabolomics of glycerolipid and glycerophospholipid metabolism in early MASLD. (B) Gene expression changes in fatty acid β-oxidation, mitochondrial respiratory chain, and lipid droplet (LD) metabolism associated with steatosis and fibrosis. There was a notable downregulation of genes involved in the electron transport chain of the inner mitochondrial membrane, specifically in complex I (NDUF family), complex II (SDH family), complex III (UQCR), complex IV (COX), and complex V (ATP5 family) as fibrosis progressed. Genes involved in respiratory electron transport were colored by the significance (–log10(p value)*sign(coefficient)) in relation to fibrosis grades. Inset: gene expression changes in the electron transport chain are depicted.

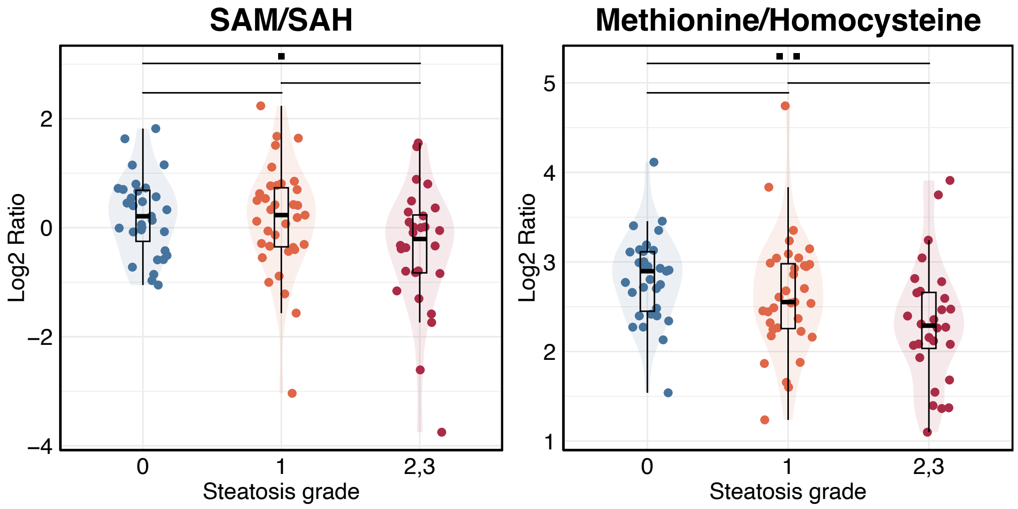


**Figure 3–figure supplement 2. Ratios of metabolites in one-carbon metabolism in individuals with different steatosis grades.** SAM, S-adenosylmethionine. SAH, S-adenosylhomocysteine.

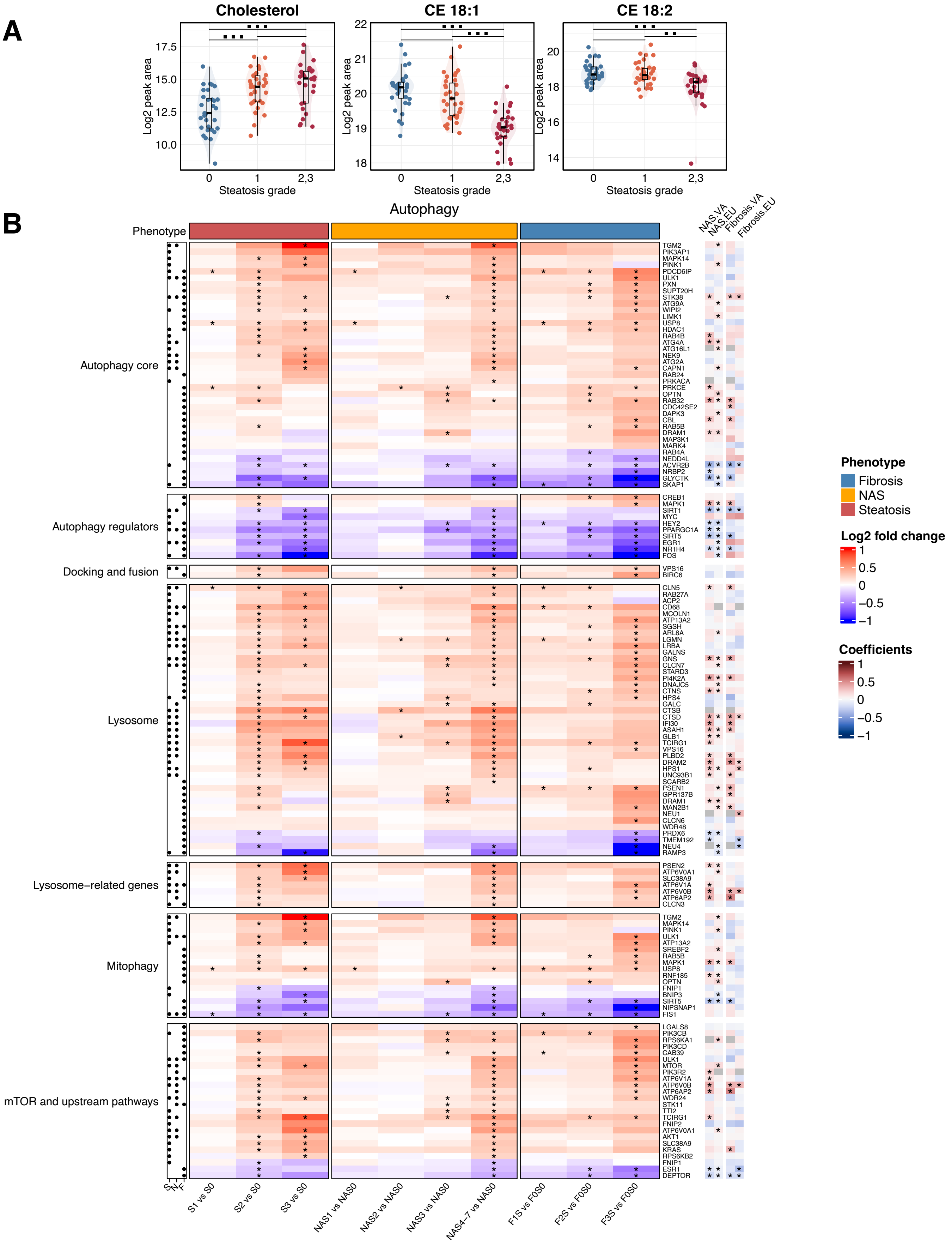


**Figure 4–figure supplement 1. Autophagy regulation in the liver.**
(A) Hepatic free cholesterol, CE 18:1, and CE 18:2 levels in this cohort in relation to steatosis grade (x-axis). (B) Heatmap of autophagy genes in relation to histological features, indicating active lipophagy in the liver of obese individuals. Asterisks in the main heatmap denote the significance ($p < 0.05$) from comparisons of each subgroup. Association with steatosis (S), NAS (N) and fibrosis (F) was indicated by black dots ($q < 0.05$, left). Results were cross-referenced with two published cohorts (VA cohort, GSE130970; EU cohort, GSE135251) (*$q < 0.05$ and with consistent direction of changes as in our cohort, bottom).

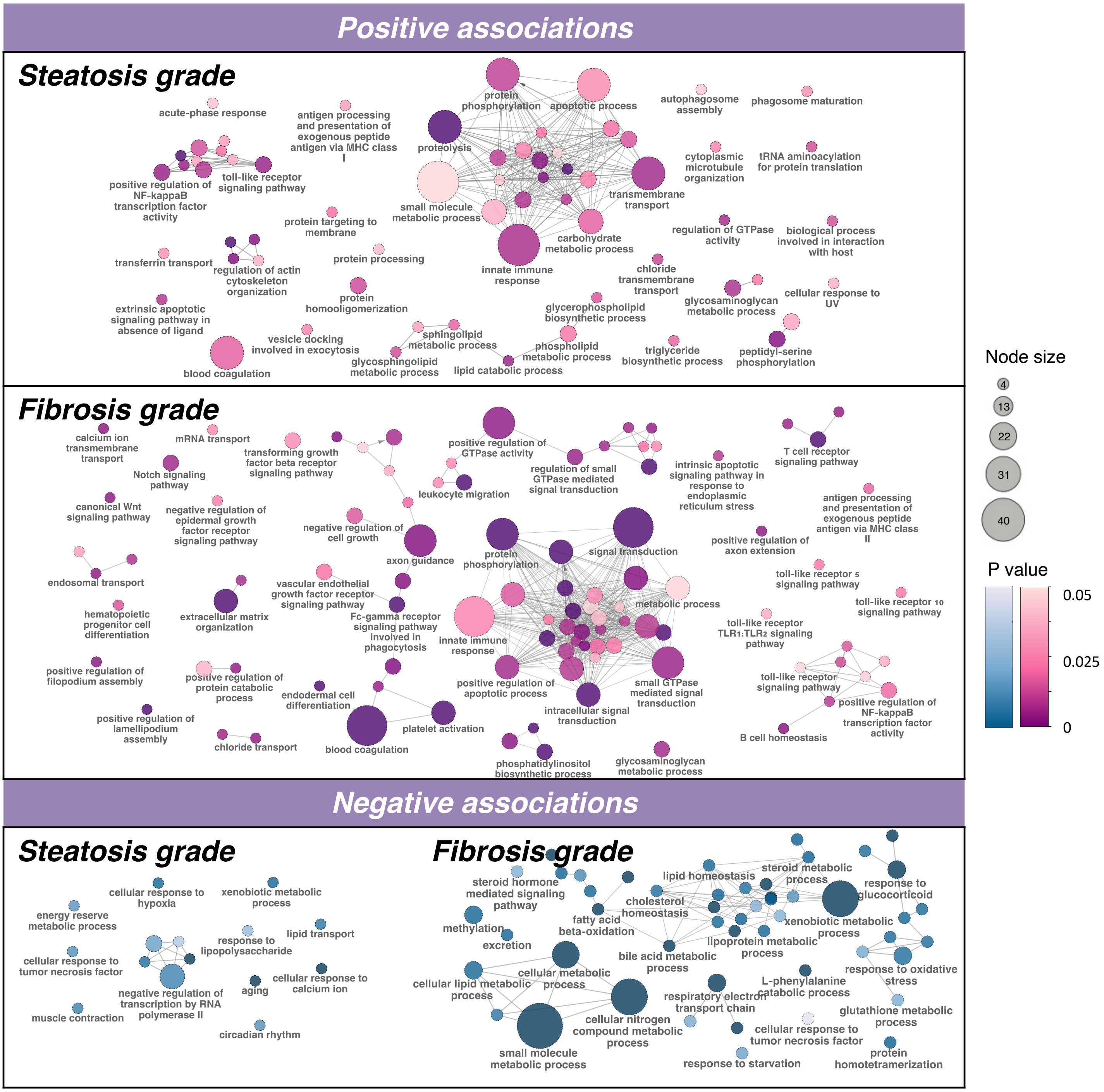


**Figure 5–figure supplement 1. Functional enrichment of gene sets associated with steatosis and fibrosis in the liver transcriptome.** Node size is depicted by circles and P value by color.

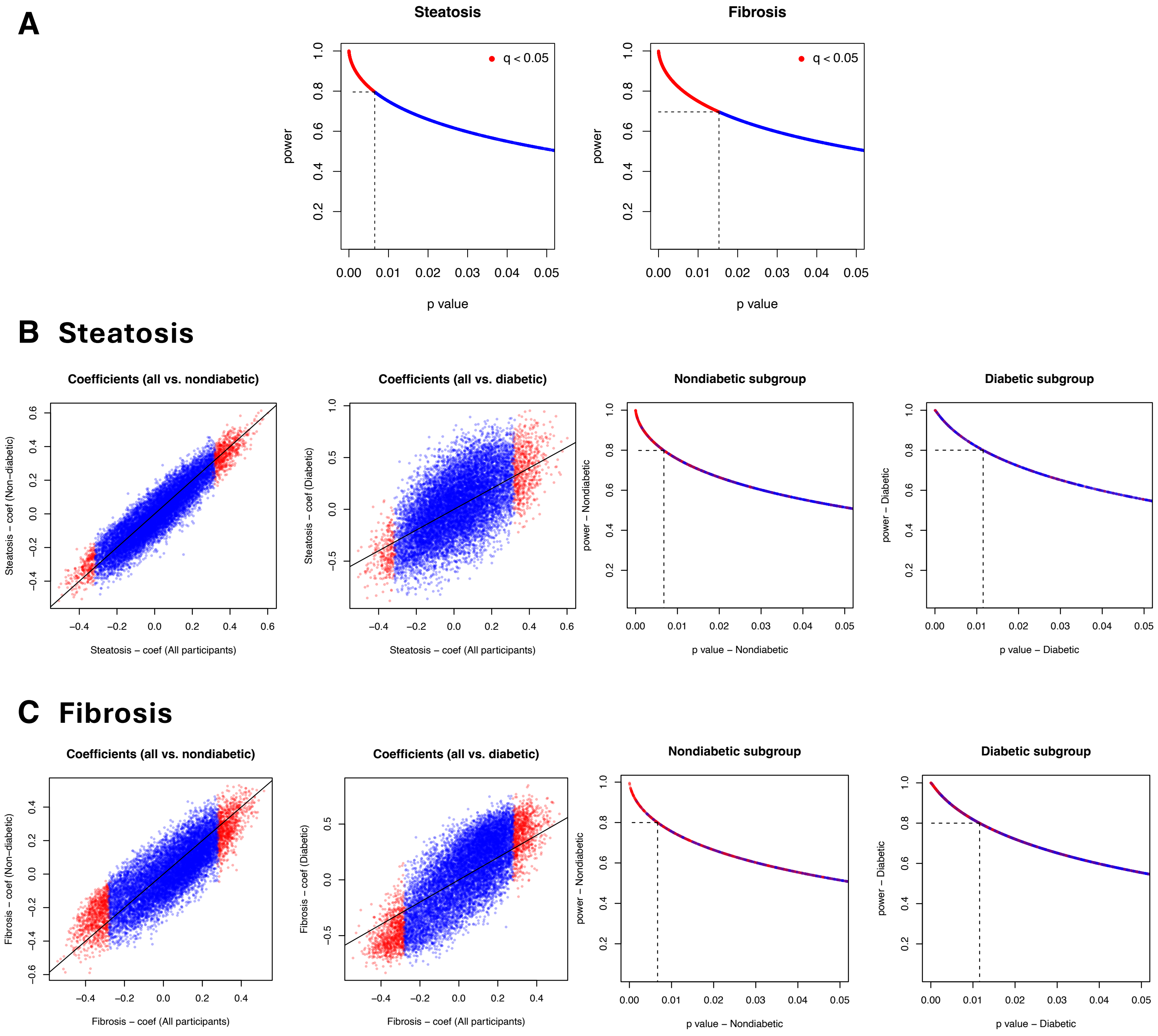


**Figure 5–figure supplement 2. Statistical power and subgroup analysis of associations between gene expressions and steatosis or fibrosis grades.**
Statistical power in the original linear regression analysis of steatosis and fibrosis was plotted against p-values (A). Differential features associated with steatosis ($q < 0.05$) showed power > 0.8, whereas those associated with fibrosis showed power > 0.7. Subgroup analyses of participants with and without diabetes were conducted to evaluate associations between gene expression and steatosis (B) and fibrosis (C). The x-axis shows coefficients from the original analysis including all participants. Corresponding coefficients from the analyses of non-diabetic (left panel) and diabetic (middle-left panel) individuals are displayed alongside the statistical power for each subgroup analysis (middle-right and right panels). Red dots represent genes that were significant in the original analysis ($q < 0.05$). Subgroup analyses suggest that the results in the non-diabetic subgroup ($n = 71$) were highly consistent with findings from the original analysis ($n = 94$, adjusted for diabetes), indicating that the originally reported gene signatures, after correction for diabetic status, remain valid in non-diabetic individuals.

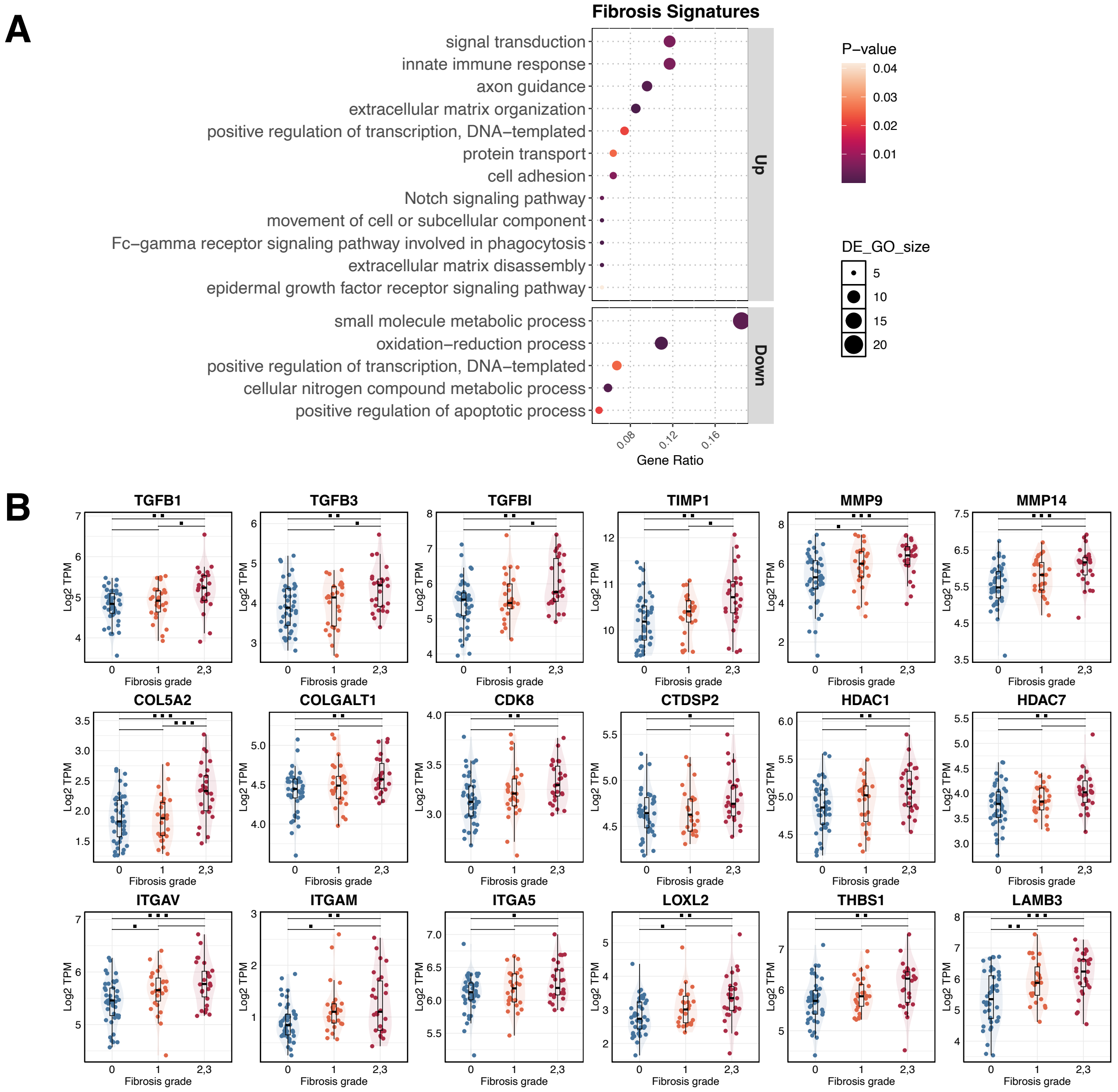


**Figure 6–figure supplement 1. Liver fibrosis pathways and gene signatures**

(A) Pathway analysis of the enrichment of pathways for the 213 fibrosis signatures. (B) TGF-β and SMAD related gene expression associated with fibrosis grades (x-axis) is depicted. Significance is indicated by black dots; * $p < 0.05$, ** $p < 0.01$, *** $p < 0.001$.

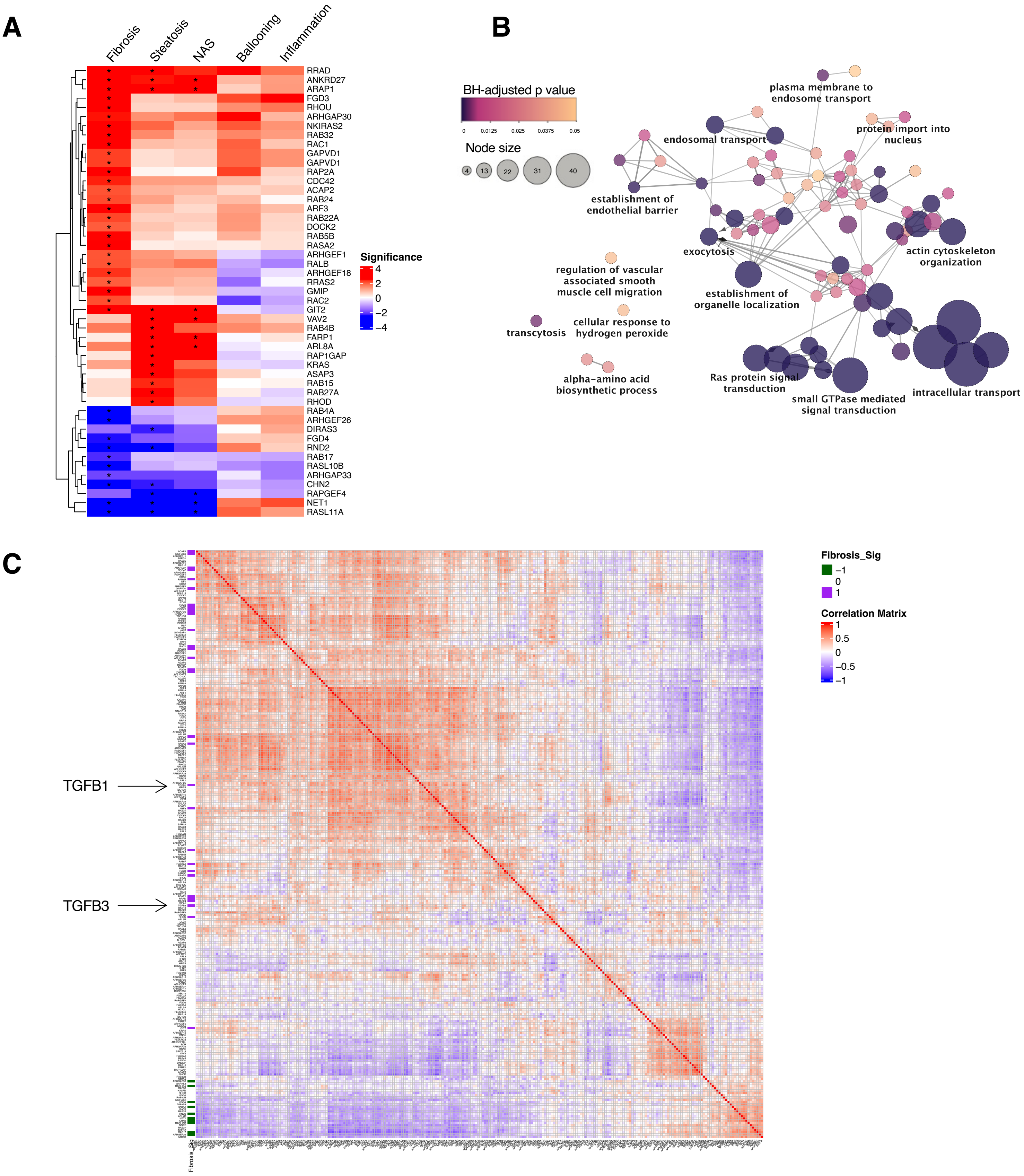


**Figure 6–figure supplement 2. GTPases and their regulation emerges as a potential target for liver fibrosis.**

(A) Heatmap of GTPase-related genes display a better correlation with fibrosis grades. Asterisks in the main heatmap denote the significance ($p < 0.05$) from comparisons of each subgroup. (B) Enrichment map of selected genes constituting the core of the protein-protein interaction network of GTPase-related genes. Nodes are colored according to BH (Benjamini-Hochberg) adjusted p-value, and node sizes are proportional to the number of genes in the term. (C) Correlation matrix of gene expression levels for TGF-β and GTPase-related genes. The correlation plot was created of Pearson's r values for GTPase-related genes and TGF-β genes (TGFB1 and TGFB3). The left annotation bar indicates the significance of associations between gene expression and fibrosis grades.

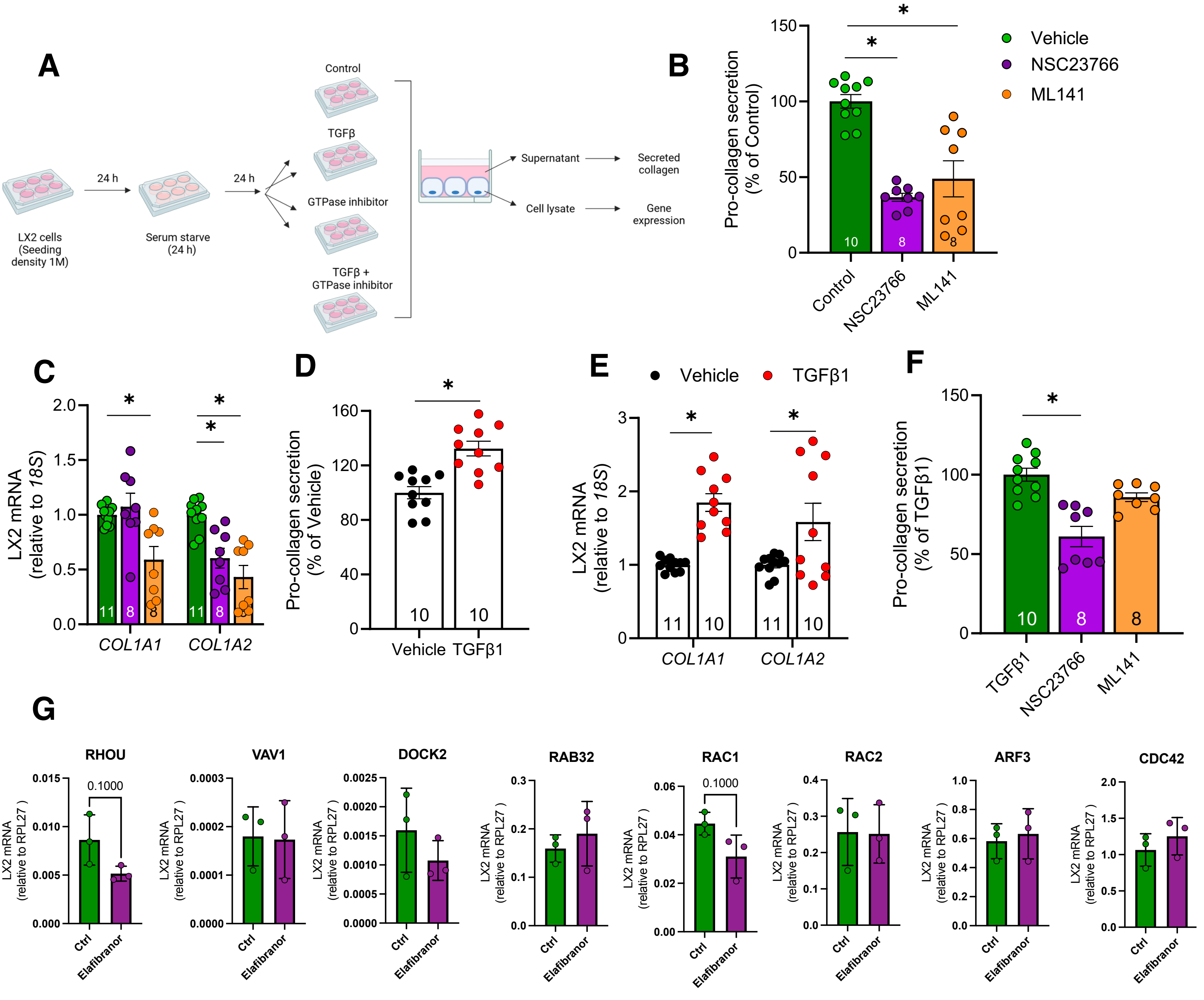


**Figure 6–figure supplement 3. GTPase inhibition and Elafibranor treatment in LX2 cells.**
(A) Workflow of the LX2 experiment in panel A-F (n = 8-10). (B) GTPase inhibitors NSC23766 (Rac1 inhibitor) and ML141 (Cdc42 inhibitor) significantly reduced pro-collagen secretion from HSC-like LX-2 cells and (C) inhibited gene expression of COL1A1 and COL1A2 under basal conditions. (D) TGFβ administration increased pro-collagen secretion by 32% and (E) collagen gene expression in LX2 cells. (F) NSC23766-mediated GTPase inhibition impairs pro-collagen secretion from LX-2 cells after TGF-β treatment. Pro-collagen secretion was determined by ELISA and gene expression levels were assessed by qPCR. Asterisks (*) denote p value < 0.05 for statistical significance from one-way ANOVA with Holms-Sidak multiple comparisons (B&C), unpaired t-test (D&E) Kruskal Wallis Test with Dunns multiple comparisons (F). (G) Gene signatures in LX-2 cells with and without Elafibranor (n = 3).

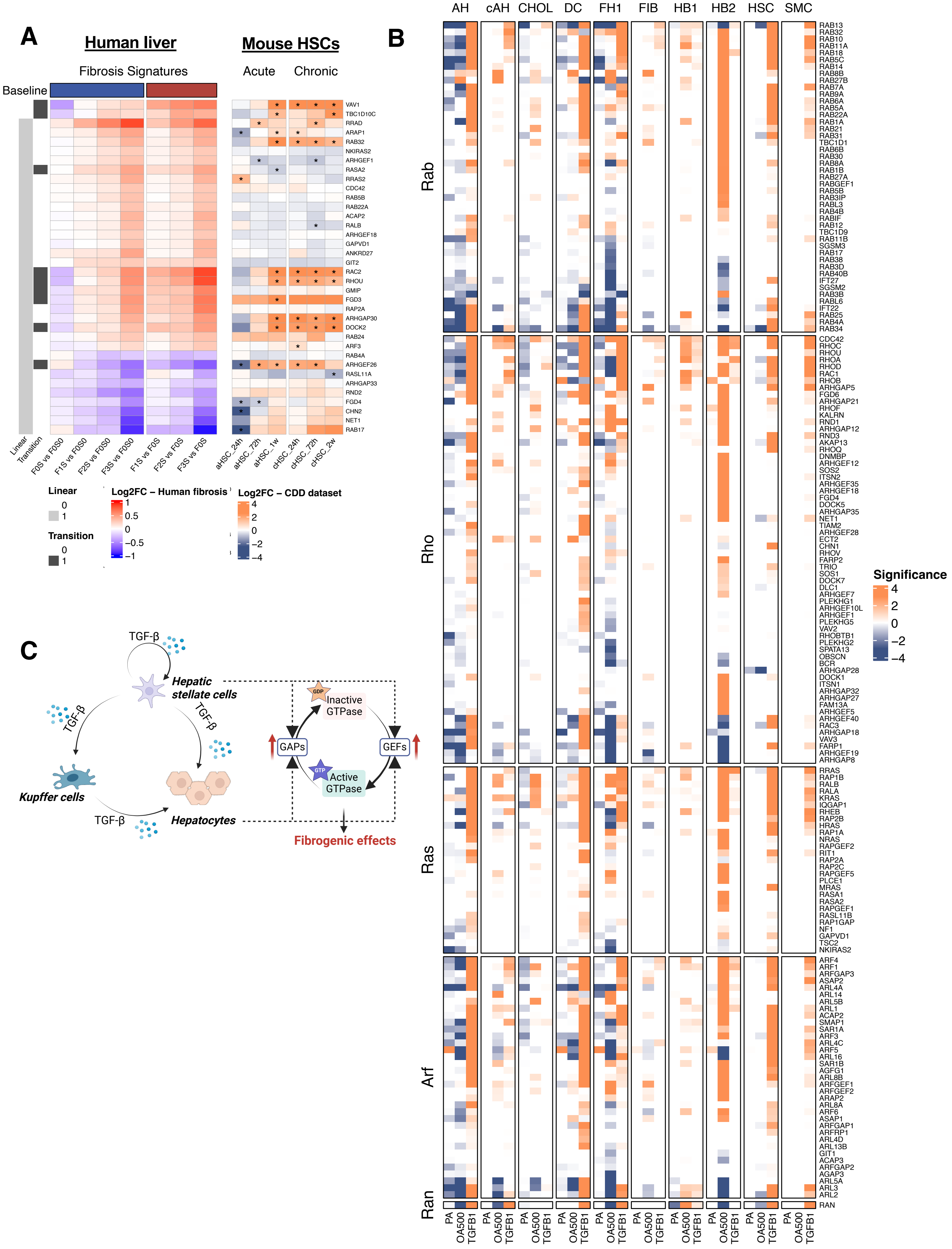
A
Human liver
Mouse HSCs
Fibrosis Signatures
Acute
Chronic
Baseline
VAV1
TBC1D10C
RRAD
ARAP1
RAB32
NKIRAS2
ARHGEF1
RASA2
RRAS2
CDC42
RAB5B
RAB22A
ACAP2
RALB
ARHGEF18
GAPVD1
ANKRD27
GIT2
RAC2
RHOU
GMIP
FGD3
RAP2A
ARHGAP30
DOCK2
RAB24
ARF3
RAB4A
ARHGEF26
RASL11A
ARHGAP33
RND2
FGD4
CHN2
NET1
RAB17
Linear
Transition
F0S vs F0S0
F1S vs F0S0
F2S vs F0S0
F3S vs F0S0
F1S vs F0S
F2S vs F0S
F3S vs F0S
aHSC_24h
aHSC_72h
aHSC_1w
cHSC_24h
cHSC_72h
cHSC_2w
Linear
0
1
Transition
0
1
Log2FC – Human fibrosis
1
0.5
0
−0.5
−1
Log2FC – CDD dataset
4
2
0
−2
−4
B
AH
cAH
CHOL
DC
FH1
FIB
HB1
HB2
HSC
SMC
Rab
Rho
Ras
Arf
Ran
RAB13
RAB32
RAB10
RAB11A
RAB18
RAB5C
RAB14
RAB8B
RAB27B
RAB7A
RAB9A
RAB6A
RAB5A
RAB22A
RAB1A
RAB21
RAB31
TBC1D1
RAB6B
RAB30
RAB8A
RAB1B
RAB27A
RABGEF1
RAB5B
RAB3IP
RABL3
RAB4B
RABIF
RAB12
TBC1D9
RAB11B
SGSM3
RAB17
RAB38
RAB3D
RAB40B
IFT27
SGSM2
RAB3B
RABL6
IFT22
RAB25
RAB4A
RAB34
CDC42
RHOC
RHOU
RHOA
RHOD
RAC1
RHOB
ARHGAP5
FGD6
ARHGAP21
RHOF
KALRN
RND1
ARHGAP12
RND3
AKAP13
RHOQ
DNMBP
ARHGEF12
SOS2
ITSN2
ARHGEF35
ARHGEF18
FGD4
DOCK5
ARHGAP35
NET1
TIAM2
ARHGEF28
ECT2
CHN1
RHOV
FARP2
TRIO
SOS1
DOCK7
DLC1
ARHGEF7
PLEKHG1
ARHGEF10L
ARHGEF1
PLEKHG5
VAV2
RHOBTB1
PLEKHG2
SPATA13
OBSCN
BCR
ARHGAP28
DOCK1
ITSN1
ARHGAP32
ARHGAP27
FAM13A
ARHGEF5
ARHGEF40
RAC3
ARHGAP18
VAV3
FARP1
ARHGEF19
ARHGAP8
RRAS
RAP1B
RALB
RALA
KRAS
IQGAP1
RHEB
RAP2B
HRAS
RAP1A
NRAS
RAPGEF2
RIT1
RAP2A
RAP2C
RAPGEF5
PLCE1
MRAS
RASA1
RASA2
RAPGEF1
RASL11B
RAP1GAP
NF1
GAPVD1
TSC2
NKIRAS2
ARF4
ARF1
ARFGAP3
ASAP2
ARL4A
ARL14
ARL5B
ARL1
ACAP2
SMAP1
SAR1A
ARF3
ARL4C
ARF5
ARL16
SAR1B
AGFG1
ARL8B
ARFGEF1
ARFGEF2
ARAP2
ARL8A
ARF6
ASAP1
ARFGAP1
ARFRP1
ARL4D
ARL13B
GIT1
ACAP3
ARFGAP2
AGAP3
ARL5A
ARL3
ARL2
RAN
PA
OA500
TGFB1
Significance
4
2
0
−2
−4
C
TGF-β
Hepatic stellate cells
TGF-β
TGF-β
Kupffer cells
TGF-β
Hepatocytes
GDP
Inactive GTPase
GAPs
GEFs
GTP
Active GTPase
Fibrogenic effects

**Figure 6–figure supplement 4**. **GTPase-related genes in external systems**
(A) Human fibrosis signatures from the present study were partially reflected in a mouse study (GSE176042), where RNA sequencing was performed on hepatic stellate cells (HSCs) isolated from both acute and chronic mouse fibrosis models. GTPase-related genes, especially those involved in the initiation of fibrosis, were also upregulated in HSCs isolated from both acute and chronic liver fibrosis mouse models at different time points. Row annotations 'Linear' and 'Transition' indicate whether gene expression is linearly associated with fibrosis grades (Linear = 1) or shows a significant difference between patients with fibrosis and steatosis versus simple steatosis (Transition = 1). Asterisk denotes genes with adjusted p value $< 0.05$. (B) GTPase-related genes induced by TGF-β were elevated in hepatocytes and HSCs at different stages of a human liver organoid system (GSE207889). Significance was calculated by –log10(p value)*sign(fold change). Asterisk denotes genes with adjusted p value $< 0.05$. AH, adult hepatocyte-like; CHOL, cholangiocyte-like; DC, ductal cell-like; FH1, fetal hepatocyte 1-like; FIB, fibroblast-like; HB1, hepatoblast 1-like; HB2, hepatoblast 2-like; HSC, hepatic stellate cell-like; SMC, smooth muscle cell-like. (C) Schematic illustration of the hypothesized role of GTPase signaling in regulating liver fibrogenesis. TGF-β production is initiated in non-parenchymal liver cells, leading to the overexpression of GTPase regulators in both hepatic stellate cells and hepatocytes. This activation of GTPase signaling amplifies downstream fibrogenic effects, creating a feed-forward loop that intensifies fibrosis progression. Further investigation is needed to prove by which mechanisms GTPases regulate collagen expression and therefore fibrosis.

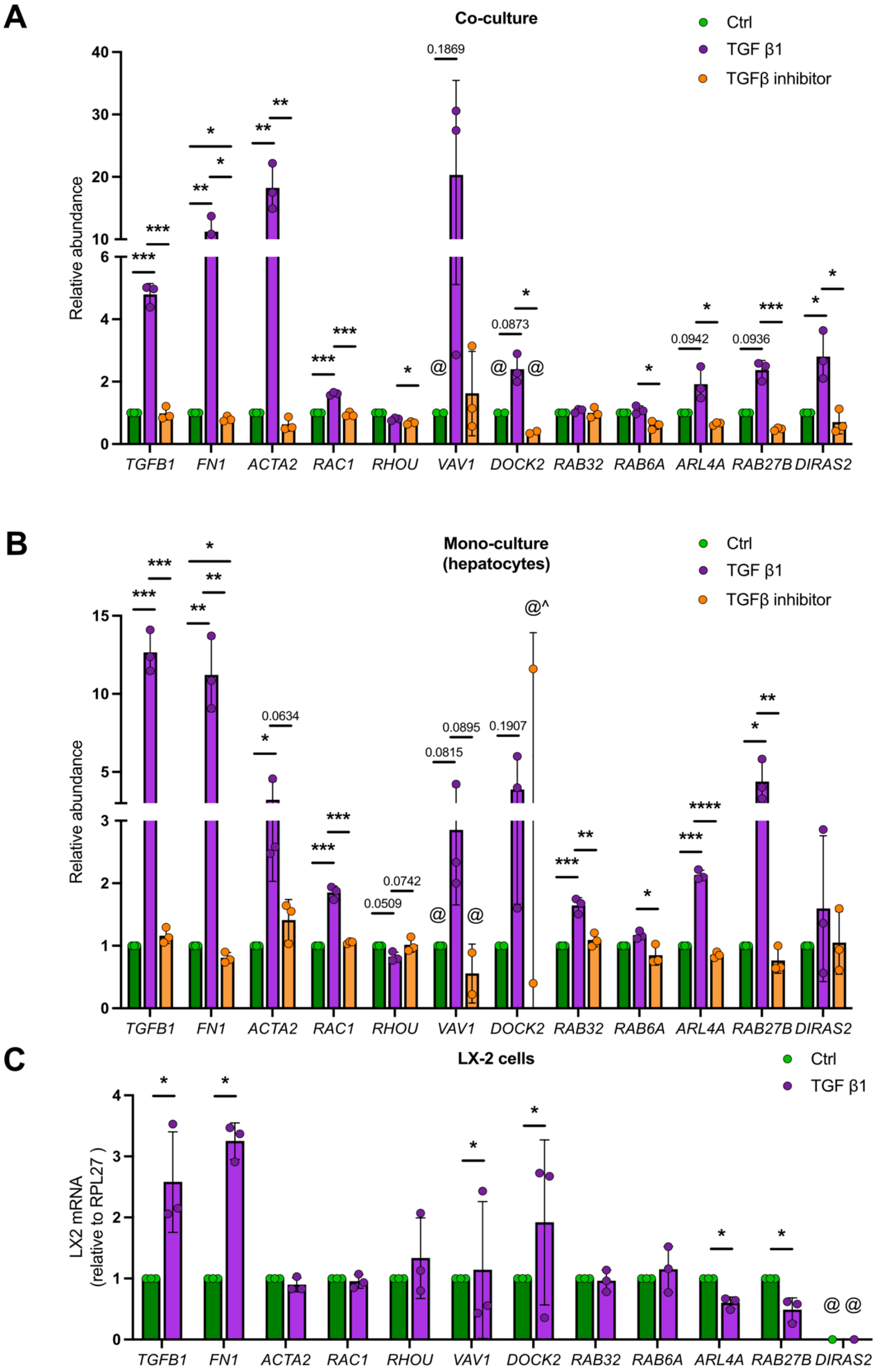


**Figure 6–figure supplement 5. Expression of GTPase-related genes in spheroid co-culture, hepatocyte monoculture, and LX-2 cells.** RNA-sequencing-based expression of GTPase-related genes in spheroid co-culture, hepatocyte monoculture upon control (Ctrl), TGF-β1 or TGFβ – inhibitor treatment (A, B). Expression of GTPase-related genes in control (Ctrl) and TGF-β1- treated (24h) LX2 cells measured by qPCR (C). Asterisks (*) denote p value < 0.05, (**) p < 0.005, (***) p < 0.0001 for statistical significance from unpaired t-test, numerical values above the bars indicate p-value. Bars were removed where data did not show strong directionality and statistical reliability, as indicated by (^) symbol. Symbol (@) indicates one or more expression values were not detected in the original dataset. Please note that ACTA2 expression in LX-2 cells is time dependent (unpublished results) and therefore we also included fibronectin (FN1) as a control for TGF-β1 treatment.
